# Serotonergic and Noradrenergic Interactions in Pupil-linked Arousal

**DOI:** 10.1101/2025.03.26.644382

**Authors:** Maxime Maheu, Tobias H. Donner, J. Simon Wiegert

## Abstract

Changes in central arousal state shape cortical computations underlying perception, thought, and action. Variations in arousal are accompanied by fluctuations in pupil size. In turn, pupil dynamics are often used as a marker of noradrenaline release from neurons of the locus coeruleus. The serotonergic system of the dorsal raphe also contributes to the brainstem control of waking states. However, little is known about its relationship to noradrenergic activity and arousal dynamics in awake animals. Here, we simultaneously accessed both systems in awake mice and unraveled their unique and joint contributions to pupil-linked arousal. Serotonergic and noradrenergic systems co-fluctuated, and serotonergic dorsal raphe neurons affected pupil size partly via noradrenergic populations in the locus coeruleus. Yet, part of the serotonergic control of pupil dynamics was independent of the locus coeruleus. Our findings challenge common assumptions about the neuromodulatory control of pupil dynamics and illuminate the interplay between distinct neurochemical systems within the arousal network of the brainstem.

## Introduction

The ascending modulatory systems of the brainstem, such as the locus coeruleus (LC) noradrenaline (NA) system or the serotonergic (5HT) system of the dorsal raphe (DR) nucleus, control the state of cortical networks, and thereby sculpt perception, thought, and action^1–10^. These systems consist of richly connected but neurochemically distinct neuronal ensembles with diffuse ascending projections to the forebrain that release different modulatory neurotransmitters in their target networks^11^. While extensive research has elucidated the contribution of these different systems to cognitive and behavioral processes^1,12–15^, these studies have mostly considered each system in isolation. Thus, a better understanding of how these neuromodulatory systems cooperate and uniquely contribute to the regulation of brain states is needed.

A key physiological readout of cortical state in awake animals is non-luminance-mediated variations in pupil size, often referred to as pupil-linked arousal^7,10^. Importantly, these concomitant changes of pupil size and cortical state occur also in periods of quiescence, in absence of any locomotion^4^. Work in rodents, monkeys, and humans has established a direct link between these pupil dynamics and the activity of the LC-NA system^1–10,16–24^. An important question is whether, and how, modulatory brainstem systems other than LC-NA affect pupil size^7,10^. While recent studies have suggested a role for the cholinergic, dopaminergic, and DR-5HT systems^22–29^, evidence is often indirect or correlative and the interplay between these other systems and LC-NA is rarely addressed. Given the rich anatomical connectivity between the different brainstem nuclei^30–37^, links between each structure and pupil size (or any other behavioral readout) are difficult to interpret without concurrently assessing activity in other nuclei.

The interplay between the LC-NA and DR-5HT systems is of particular interest given their complex and sometimes opposing effects on arousal, sleep^18,38–41^, sensory processing^42–44^, and high-level cognitive functions^14,45^. Critically, both the LC-NA and DR-5HT systems are implicated in several neuropsychiatric disorders^14^ and major targets in the pharmacological treatment of depression^11^. Therefore, detailed investigations of the interplay between LC-NA and DR-5HT systems in awake animals is required to understand the natural control of brain states as well as ultimately aberrations thereof in disease.

Here, we set out to systematically identify the causal functional relationships between DR-5HT, LC-NA, and pupil-linked arousal in awake mice. To do so, we leveraged recently developed genetic tools to simultaneously monitor (with calcium indicators) and manipulate (with optogenetic actuators) both regions, while tracking pupil dynamics and occasional periods of locomotion. We tested four hypothetical scenarios describing the putative regulation of pupil dynamics by DR-5HT and the involvement of LC-NA in this process (Fig. 1a): DR-5HT does not regulate pupil size (*H*_0_); DR-5HT regulates pupil size in an LC-NA-independent manner (*H*_1_); DR-5HT regulates pupil size in an LC-NA-dependent manner (*H*_2_); or a mixture of the two previous scenarios whereby both pathways would co-exist (*H*_3_). Note that these functional architectures could be realized in multi-synaptic pathways also including nuclei other than LC and DR (Supplementary Fig. 1).

**Fig. 1.**
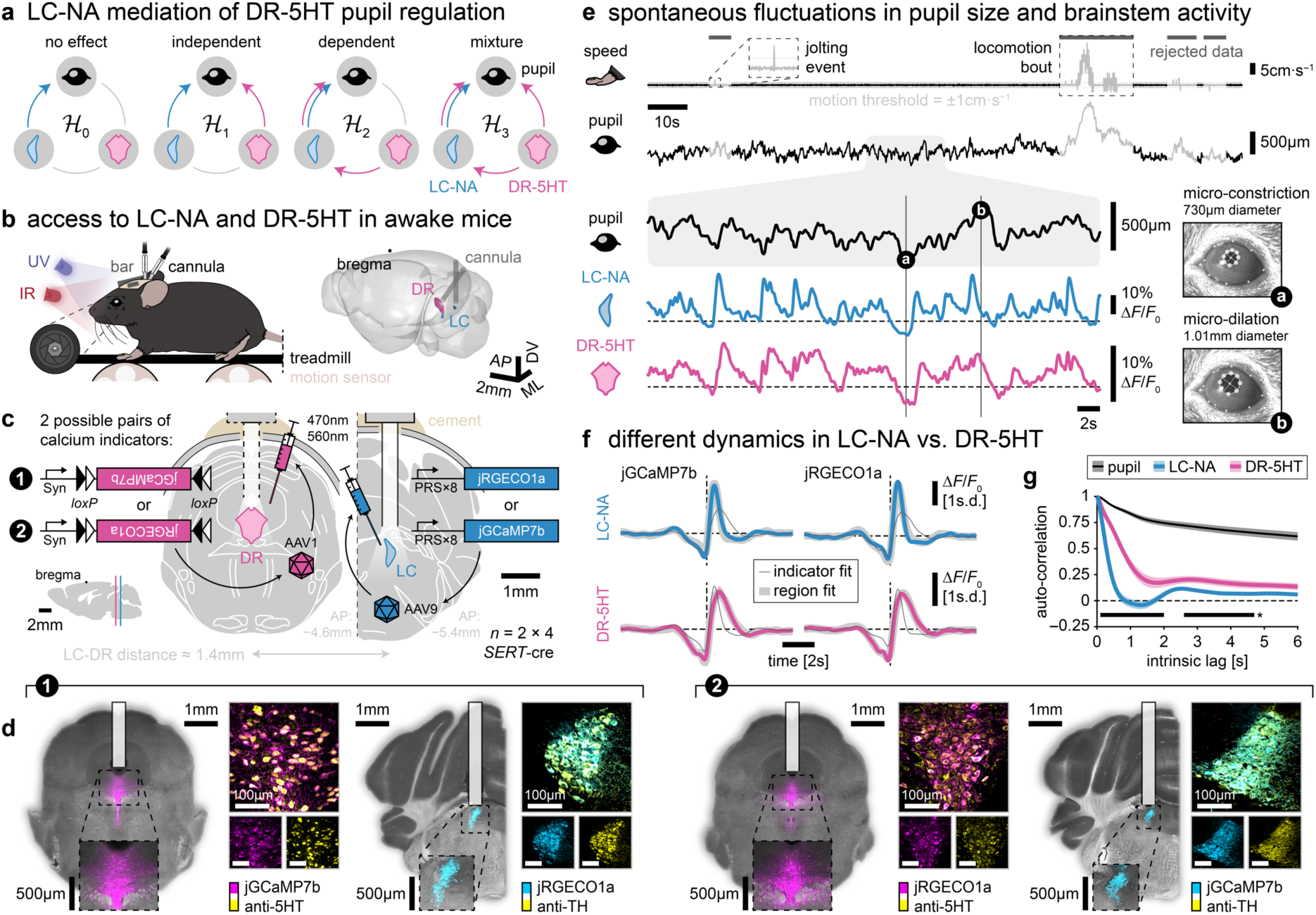
Spontaneous dynamics of pupil size, LC-NA and DR-5HT activities in mice. **a,** Alternative hypotheses for DR-5HT control of pupil and its regulation by LC-NA. Arrows depict functional modulation, not anatomical connections. **b,** Left: schematic of the experimental apparatus where a head-fixed mouse (adapted from mindthegraph.com) can walk on a linear treadmill while instantaneous speed is recorded, and the left eye is video monitored by an infrared (IR) camera. Ambient light is controlled and maintained constant by an ultraviolet (UV) light directed to the non-recorded eye at a power level maximizing the pupil dynamic range. Right schematic: LC, DR and corresponding implanted fiber optics shown within a mouse glass brain from the 10μm-resolution *Allen Mouse Brain Common Coordinate Framework* atlas^57^. The LC is inflated to double its size for visualization purposes. The DR is accessed through a 32° posterior angled approach. **c,** Representative coronal brainstem slices from the *Paxinos Stereotaxic Atlas*^58^ showing LC and DR. LC and DR are injected with one of two spectrally distinct calcium indicators (jGCaMP7b or jRGECO1a) in a counter-balanced manner (4 mice with LC^jGCaMP7b^-DR^jRGECO1a^, 4 mice with LC^jRGECO1a^-DR^jGCaMP7b^). Expression was restricted to NA cells in the LC by using the synthetic PRS×8 promoter and to 5HT cells in the DR using cre-dependent constructs and a transgenic mouse line (*SERT*-cre). Excitation and emission light was delivered and collected through a fiber optic implant. **d,** Confirmation of cell-type-specific expression of calcium indicators in LC-NA/DR-5HT by histology and immunostaining against TH/5HT (different brain slices in left and right views). **e,** Snapshot of instantaneous speed, pupil size, LC-NA and DR-5HT activity. Gray lines: motion-related sections of rejected data, defined as samples within a window (−3 and +8.5s) around samples with speed ≥ 1cm·s^−1^, capturing both locomotion bouts and isolated jolting events. Eight pupil markers, as identified by a deep neural network^59^, are overlaid on eye frames corresponding to micro-constriction/-dilation events. Pupil size corresponds to the diameter of the fitted ellipse. **f,** Mean brainstem activity transients (5% most extreme first-order derivative, 400ms minimum duration and non-overlapping transients) for each combination of brainstem region and calcium indicator. Thick light grey/thin dark grey line: fitted activity transient from the same brainstem region/calcium indicator. **g,** Mean ± standard error of the mean (s.e.m.) auto-correlogram for LC-NA, DR-5HT and pupil. Horizontal black lines: significant difference between LC-NA and DR-5HT (cluster-based permutation test, *p* < 0.05).

We discovered a cooperative interaction between DR-5HT and LC-NA activity, through both correlation of their natural, spontaneous dynamics during wakefulness, as well as through simultaneous activation of DR-5HT and recording of LC-NA activity. DR-5HT activity translated into pupil dilation, and part of this effect was mediated by LC-NA activity. Crucially, however, we also identify an LC-NA-independent contribution of DR-5HT to pupil dilation.

## Results

To explore the different functional architectures sketched in Fig. 1a, we first recorded the spontaneous activity of populations of NA neurons in the LC and 5HT neuronal populations in the DR, and related each to pupil size. We reasoned that the existence of anatomical connections between DR and LC^30–37^ (see *H*_2_ and *H*_3_) should produce co-fluctuations in activity that only concurrent recordings of the activity of both nuclei could identify. We thus established a multiplexed genetic strategy to target both neuromodulatory populations in the same mice. A synthetic promoter (PRS×8) restricted transgene expression to NA cells^46,47^ and a transgenic mouse line (*SERT*-cre) restricted the expression of cre-dependent constructs to 5HT cells (Fig. 1b-d)^48,49^.

Given their close proximity within the brainstem (∼1.4mm), we targeted LC with a straight approach and DR with a posterior angled approach (Fig. 1b; as previously validated^49,50^), and used spectrally distinct, genetically encoded calcium indicators (jGCaMP7b and jRGECO1a) to monitor their activity simultaneously. These indicators were conditionally expressed in LC-NA vs. DR-5HT in a counter-balanced way to control for the different biophysical properties of the indicators (dynamic range, kinetics, etc.). Possible cross-talk between the two indicators was avoided by a tailored set of emission and excitation filters, as well as time-division multiplexed acquisition (Supplementary Fig. 2). In addition, we confirmed the absence of expression in the non-targeted region (Supplementary Fig. 3) and we verified reliable fiber positioning close to the targeted brainstem regions (Supplementary Fig. 4). Finally, to rule out the possibility that differences between brainstem structures would be due to asymmetric signal quality, we confirmed that signal-to-noise (measured as Δ*F*/*F*_0_) was comparable across brainstem regions and calcium indicators at the group level (Supplementary Fig. 5).

### Spontaneous dynamics of brainstem activity

Spontaneous fluctuations of LC-NA and DR-5HT activity as well as pupil size were recorded from passive, head-fixed mice that were allowed to walk along a linear treadmill while their left eye was video monitored (Fig. 1b). Dilations and constrictions of the pupil were preceded by increases and decreases, respectively, in both LC-NA and DR-5HT activity (Fig. 1e).

In line with previous studies^5,22,28,51,52^, locomotion was accompanied by pupil dilation, starting before locomotion onset (∼3s) and ending long (∼8.5s) after locomotion offset. Given that locomotion alters brain-wide activity in mice^53^ and creates locomotion-related artefacts in recordings that are difficult to correct for^54–56^, we focused all further analyses unless stated otherwise on periods of quiescence, excluding locomotion bouts (see Methods).

Population dynamics were faster for LC-NA than DR-5HT. Individual calcium transients were wider in DR-5HT compared to LC-NA (Fig. 1f), an effect that was independent of the calcium indicator (Supplementary Fig. 6). Likewise, the auto-correlogram was steeper for LC-NA than DR-5HT (Fig. 1g), again regardless of calcium indicator (Supplementary Fig. 7). Given the negligible impact of calcium indicator, we pooled population dynamics obtained across the two combinations of calcium indicators for all subsequent analyses.

### Co-fluctuations of LC-NA and DR-5HT

The activity of the two brainstem centers tended to co-fluctuate. Plotting DR-5HT activity as a function of binned LC-NA activity and DR-LC lag revealed that strong LC-NA activity levels were followed by increase in DR-5HT activity (Fig. 2a). Correspondingly, the cross-correlogram exhibited a peak at around 450ms, with LC-NA activity preceding DR-5HT activity changes (Fig. 2b-c), consistent with a predominant direction from LC-NA to DR-5HT. While the coupling between LC-NA and DR-5HT is strong at the best lag — likely reflecting LC-DR direct and indirect connectivity^30,34^ — the majority of the variance (coefficient of determination: 0.54^2^ = 29%) is still unique to each brainstem structure, leaving ample room for region-specific contributions to pupil dynamics.

**Fig. 2.**
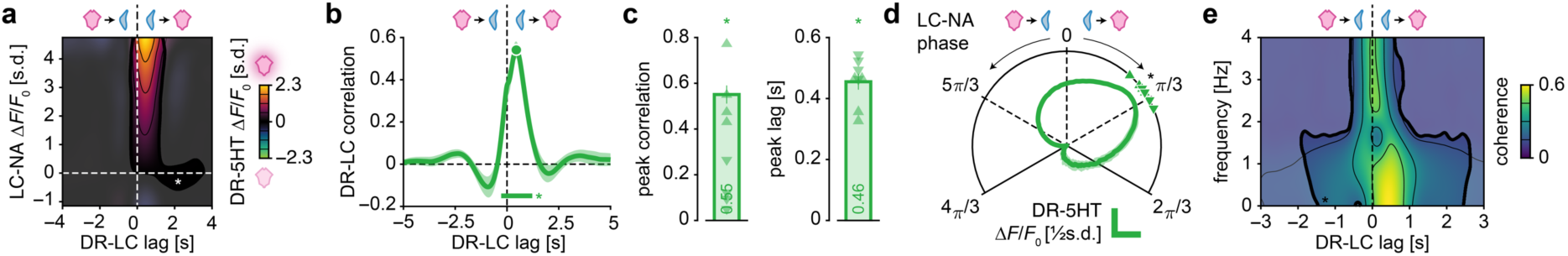
Co-fluctuations of LC-NA and DR-5HT. **a,** Mean DR-5HT activity as a function of DR-LC lag and LC-NA activity. Positive lags: LC-NA leads DR-5HT. Opaque regions: DR-5HT activity significantly different from zero (cluster-based permutation tests, *P* < 0.05, *; n.s.: non-significant). **b,** Mean ± s.e.m. cross correlogram between LC-NA and DR-5HT. Positive lags: LC-NA leading DR-5HT. Horizontal bar: correlation significantly larger than zero (cluster-based permutation test, *P* < 0.05, *). **c,** Mean ± s.e.m. DR-LC peak correlation and corresponding lag (dot in **b**). Upward-/downward-pointing triangles: individual mice expressing LC^jGCaMP7b^-DR^jRGECO1a^/LC^jRGECO1a^-DR^jGCaMP7b^. DR-LC lag significantly different from zero (Wilcoxon signed rank test, *P* < 0.05, *). **d,** Mean ± s.e.m. DR-5HT activity aligned on the Hilbert transform of band-passed LC-NA activity (0.1-1Hz). Upward-/downward-pointing triangles: individual mice expressing LC^jGCaMP7b^-DR^jRGECO1a^/LC^jRGECO1a^-DR^jGCaMP7b^. Peak of the phase-aligned DR-5HT activity significantly different from zero (Wilcoxon rank sum test, *P* < 0.05, *). **e,** Mean coherence between LC-NA and DR-5HT as a function of DR-LC lag. Positive lags: LC-NA leads DR-5HT. Opaque regions: coherence significantly different from zero (cluster-based permutation tests, *P* < 0.05, *; n.s.: non-significant).

The co-fluctuation between LC-NA and DR-5HT was, for the lower-frequency activity component (0.1-1Hz), approximately periodic with a phase shift between LC-NA (lead) and DR-5HT of about π/3 (Fig. 2d-e). In addition to this low frequency (< 1Hz) component (that dominated the broadband signal analyses in Fig. 2a-b) coherence analysis identified a separatable higher-frequency (> 2Hz) component of the co-fluctuation (Fig. 2e: in which the two brainstem nuclei were almost synchronously active. These analyses show that, LC-NA and DR-5HT co-fluctuate significantly in awake mice, with a systematic lead of LC-NA.

### Timing, strength, and dynamics of pupil-predictive activity in LC-NA vs. DR-5HT

In all above scenarios except *H*_0_, DR-5HT would exhibit pupil-predictive activity. To first reject *H*_0_, we tested if changes in DR-5HT activity would be consistently followed by changes in pupil size, as observed in the snapshot of data shown in Fig. 1. We aligned brainstem activity to fast and large pupil dilation and constriction events (Fig. 3a). These events were preceded by increases and decreases, respectively, in the activity of both LC-NA and DR-5HT. Changes in LC-NA activity seemed to precede changes in DR-5HT for both pupil constrictions and dilations. To assess to which extent this effect would generalize to the whole range of pupil fluctuations observed, we plotted brainstem activity as a function of binned pupil size and brainstem-pupil lag (Fig. 3b). The resulting maps revealed strong and weak activity clusters for large and small pupil size, respectively, in both LC-NA and DR-5HT. Activity clusters seemed stronger and temporally more extended in DR-5HT than in LC. The activity of both brainstem systems scaled monotonically with pupil size.

**Fig. 3.**
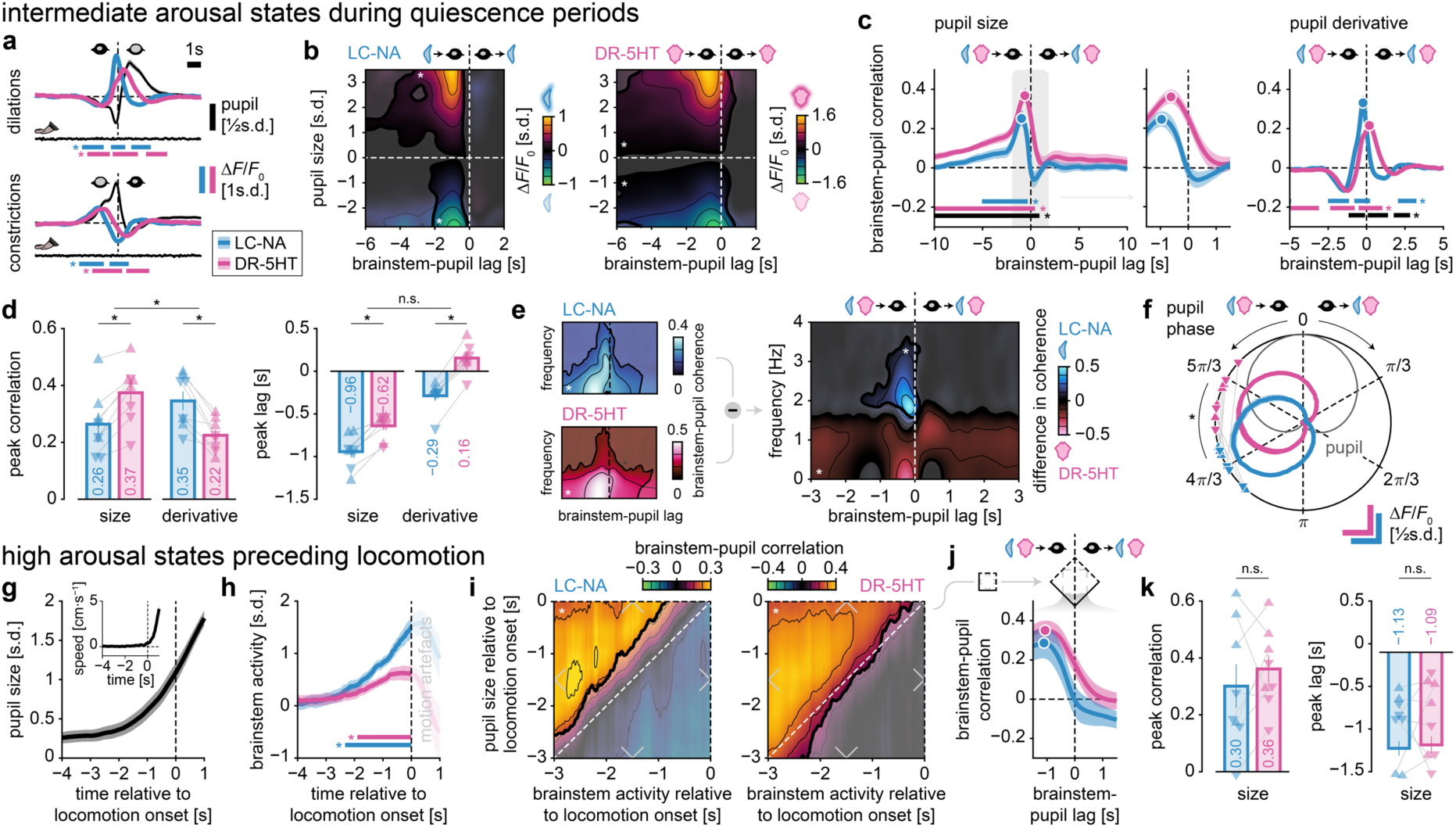
Timing, strength and dynamics of pupil-predictive activity in LC-NA vs. DR-5HT. **a,** Mean ± s.e.m. brainstem activity aligned to large and fast (5% most extreme of pupil derivative, minimum duration of 500ms) pupil dilation (top) and constriction (bottom) events. Horizontal colored bars: activity significantly different from zero (cluster-based permutation tests, *P* < 0.05, *). **b,** Mean brainstem activity as a function of brainstem-pupil lag and pupil size. Negative lags: brainstem leads pupil. Opaque regions: brainstem activity significantly different from zero (cluster-based permutation tests, *P* < 0.05, *; n.s.: non-significant). **c,** Mean ± s.e.m. brainstem-pupil cross-correlogram using pupil size (left and middle plots) or first-order derivative of pupil size (right plot). Negative lags: brainstem leads pupil. Horizontal colored bars: correlation significantly different from zero; black horizontal bars: significant difference in correlation between LC-NA vs. DR-5HT (cluster-based permutation tests, *P* < 0.05, *). **d,** Mean ± s.e.m. peak cross-correlation and corresponding brainstem-pupil lag (dots in **c**). Upward-/downward-pointing triangles: individual mice expressing LC^jGCaMP7b^-DR^jRGECO1a^/LC^jRGECO1a^-DR^jGCaMP7b^. Significant/non-significant differences between LC-NA and DR-5HT (Wilcoxon signed rank test, *P* < 0.05, *; n.s.: non-significant). **e,** Mean coherence between brainstem activity and pupil size (left maps) and difference between coherence maps (after normalization to 1; right map), both as a function of brainstem-pupil lag. Positive/negative coherence: larger/smaller for LC-NA than DR-5HT. Negative lags: brainstem leads pupil. Opaque regions: difference in coherence significantly different from zero (cluster-based permutation tests, *P* < 0.05, *; n.s.: non-significant). **f,** Mean ± s.e.m. brainstem activity aligned on a canonical cycle of pupil dilation and constriction derived from the Hilbert transform of band-passed pupil size (0.1-1Hz). Upward-/downward-pointing triangles: individual mice expressing LC^jGCaMP7b^-DR^jRGECO1a^/LC^jRGECO1a^-DR^jGCaMP7b^. Significant difference between peak of LC-NA and DR-5HT aligned on pupil phase (Wilcoxon signed rank test, *P* < 0.05, *). **g**, Mean ± s.e.m. pupil size locked on locomotion onset. Inset: locomotion episodes correspond to periods where instantaneous speed is larger than 1cm·s^−1^ (computed over a 5s moving window) for a minimum of 1s. **h**, Mean ± s.e.m. LC-NA and DR-5HT activities locked on locomotion onset. Horizontal colored bars: significantly larger than zero activity (cluster-based permutation tests, *P* < 0.05, *). Shaded area indicate brainstem activity recorded during locomotion that is likely contaminated by motion artefacts, hence rejected. **i**, Mean correlation (across locomotion episodes) between pupil size and LC-NA (left heatmap) or DR-5HT (right heatmap) activities. Opaque regions: correlation significantly different from zero (cluster-based permutation tests, *P* < 0.05, *; n.s.: non-significant). **j**, Mean ± s.e.m. brainstem-pupil cross-correlogram (same as middle plot in **c**) in periods preceding locomotion (corresponding to diagonal-centred squares in **i**). **k**, Mean ± s.e.m. peak cross-correlation and corresponding brainstem-pupil lag (dots in **j**; to compare with **d**).

We computed cross-correlograms between brainstem activity and pupil size to comprehensively quantify the timing and strength of pupil-predictive activity^22,60^ (Fig. 3c). Activity in LC-NA and DR-5HT both predicted subsequent pupil size. While this pupil-predictive activity occurred around 5s earlier for DR-5HT than for LC-NA, the *peak* in pupil-predictive activity was about 340ms earlier for LC-NA than for DR-5HT. This is consistent with a predominant portion of the drive of pupil dilation from LC-NA. The strength of pupil-predictive activity was larger for DR-5HT than for LC-NA

(coefficient of determination: 0.37^2^ = 14% vs. 0.26^2^ = 7%; Fig. 3d). Because this difference might be explained by the overall slower dynamics of the DR-5HT population (Fig. 1), we replaced the continuous population activity by discrete calcium events before computing cross-correlograms. This resulted in statistically indistinguishable pupil-predictive activity for LC-NA and DR-5HT (Supplementary Fig. 8). Similar conclusions were achieved when artificially matching LC-NA and DR-5HT auto-correlation profiles, suggesting that slow dynamics average out the jittered brainstem-pupil coupling, thereby improving apparent brainstem-pupil correlation.

Evidence suggests that rapid changes in pupil size are more specific to NA release than the slow changes in pupil size^22^. We, therefore, also computed cross-correlograms using the first temporal derivative of pupil size that captured the higher-frequency component of the pupil dynamics (Fig. 3c). Indeed, here pupil-predictive activity was stronger for LC-NA than for DR-5HT (coefficient of determination: 0.35^2^ = 12% vs. 0.22^2^ = 5%; Fig. 3d). Furthermore, a time-resolved analysis of coherence between brainstem activity and pupil size showed that the higher-frequency (∼2Hz) pupil dynamics were specifically related to LC-NA activity, while the lower frequency (< 1.5Hz) dynamics were primarily reflected in DR-5HT activity, and LC-NA to a smaller extent (Fig. 3e). Taken together, these results suggest that LC-NA is more predictive of rapid pupil dynamics while DR-5HT is (slightly) more predictive of slower pupil dynamics. This is in line with conclusions drawn for the comparison of noradrenergic versus cholinergic activity^22^ and have implications for pupillometric studies which could focus specifically on one or the other component of pupil dynamics^61–63^ (see Discussion).

An assessment of the lags between brainstem and pupil activity provided a complementary picture: the peak in pupil-predictive activity occurred earlier for LC-NA than for DR-5HT regardless of the metric of pupil dynamics used (respectively 960ms vs. 620ms before for pupil size, and 290ms before vs. 160ms after for pupil derivative; Fig. 3d). Similarly, when aligning brainstem activity onto the phase of pupil oscillations, we found that LC-NA was more anti-phased from the pupil than DR-5HT (−π/3 vs. −2·π/3; Fig. 3f). These results are consistent with the notion of a shorter synaptic pathway from LC-NA than from DR-5HT to pupil.

The previous analyses focused on spontaneous arousal dynamics in the absence of overt behavior. All animals occasionally started running on the treadmill. Such periods of locomotion are known to be preceded by a strong increase in markers of cortical state as well as pupil size^22,51,64^. We wondered whether such pupil dilations before the onset of locomotion behaviors follow similar increases in LC-NA and DR-5HT activity as we found for spontaneous pupil dilations. As expected from previous work^22,51,64^, pupil size increased strongly before locomotion onset in our data, with the pupil starting to dilate about 3s before crossing of the speed threshold (Fig. 3g). These locomotion-related dilations were accompanied by ramping activity of both the LC-NA and DR-5HT systems (Fig. 3h).

We next asked whether timing and correlations between the brainstem structures and locomotion-preceding pupil dilation were also preserved. For both brainstem regions, we found robust pupil-predictive activity locked to, and preceding, locomotion onset (coefficient of determination for LC: 0.30^2^ = 9%; and DR: 0.36^2^ = 13%; Fig. 3i-k).

The difference in strength (DR > LC) and timing (LC before DR) of pupil-predictive activity were qualitatively preserved at the group level but reduced and did not reach statistical significance anymore. The coupling between LC-NA and DR-5HT was preserved in these periods (Supplementary Fig. 9). In sum, the covariations of LC-NA, DR-5HT and pupil dynamics observed in periods of quiescence was preserved in the states of heightened arousal that lead up to overt behavior (i.e., locomotion), suggesting a general role of the interplay between LC-NA and DR-5HT activity in pupil-linked arousal.

### LC-NA and DR-5HT uniquely contribute to pupil fluctuations

The above-described correlations between pupil size, LC-NA, and DR-5HT activities (Supplementary Fig. 10) could also occur if DR-5HT affected pupil size via the LC-NA system^36,65–67^ (*H*_2_). Alternatively, a third region could drive both LC-NA and DR-5HT^30,68,69^, with only LC-NA regulating the peripheral pupil-control system. We thus aimed to test for a *unique* contribution of DR-5HT to pupil dynamics. Concurrently recording from both structures enabled us to inspect how they combine and possibly uniquely shape pupil dynamics. Importantly, all further analyses were conducted in a symmetrical manner: they can address not only LC-mediated pupil regulation by DR-5HT, but also DR-mediated pupil regulation by LC-NA.

First, to control for the correlated activities between LC-NA and DR-5HT, we assessed whether one brainstem system could still parametrically predicted pupil size under predetermined, constant activity levels of the other brainstem system, a case of stratified analysis. When holding LC-NA activity levels constant, we could still see changes in pupil size according to DR-5HT activity; this was true conversely for LC-NA (Fig. 4a). These findings generalized across a range of activity levels while brainstem-pupil coupling was reduced mostly for intermediate activity levels in both structures (Fig. 4b; U-shaped relationship).

**Fig. 4.**
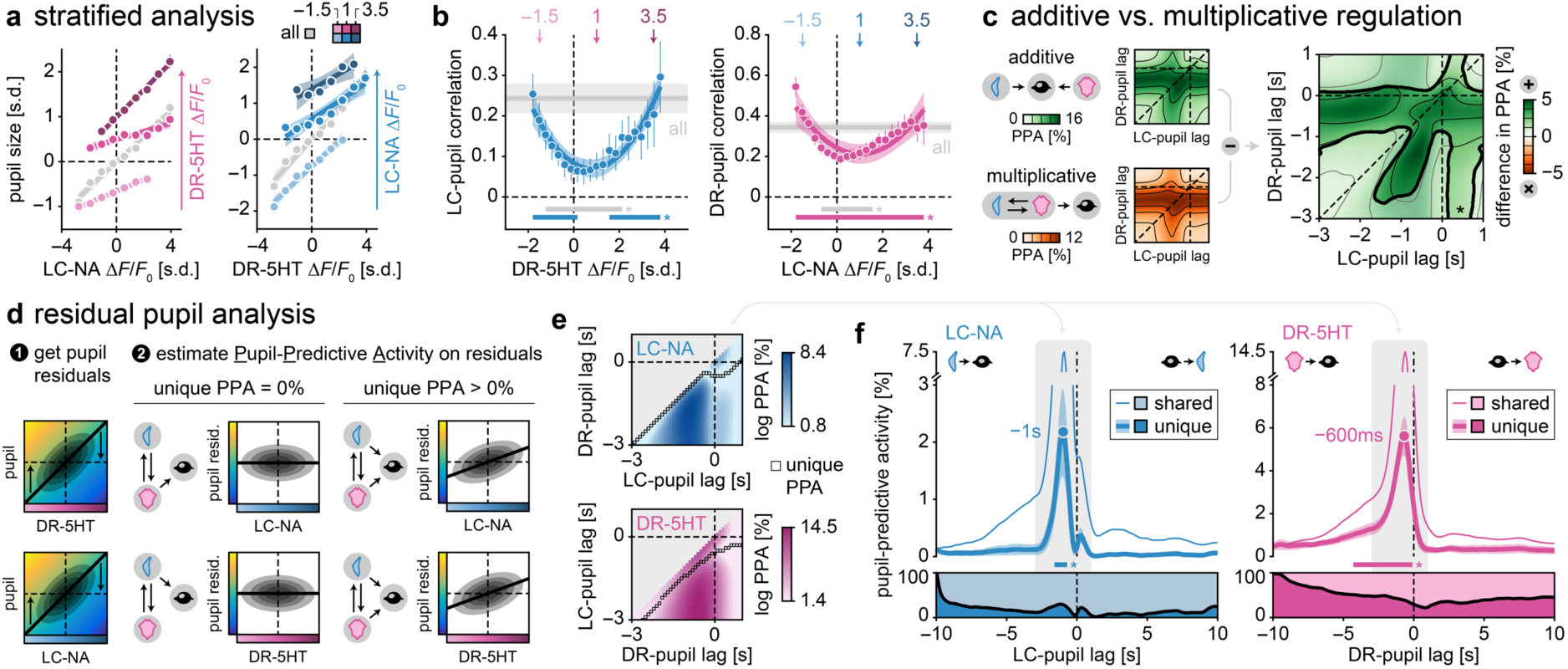
LC-NA and DR-5HT uniquely contribute to pupil fluctuations. **a,** Mean ± s.e.m. pupil size as a function of LC/DR activity while holding activity of the respective other brainstem region constant (3 levels, ±0.025s.d.), and linear fit ± 95% confidence intervals. **b,** Mean ± s.e.m. LC/DR-pupil correlation under parametric variation of activity levels of the respective other brainstem region (±0.025s.d.) and quadratic fit ±95% confidence intervals. Horizontal colored/grey bars: correlation significantly different from zero/overall correlation (i.e., without holding activity from second brainstem region; cluster-based permutation tests, *P* < 0.05, *). Arrows: example activity levels in **a**. **c,** Pupil-predictive activity (PPA) from a linear regression model where both LC-NA and DR-5HT activities combine additively (top map) or multiplicatively (bottom map) in the explanation of pupil fluctuations, and difference between the two models (right map). Opaque region: PPA significantly different between models (cluster-based permutation tests, *P* < 0.05, *; n.s.: non-significant). **e,** To estimate the unique pupil-predictive activity (PPA) in LC (top row) or DR (bottom row), we first regressed-out the contribution of DR (top) or LC (bottom) to pupil dynamics respectively. Unique PPA corresponds to the variance explained of the regression relating pupil residuals and brainstem activity. **f,** Mean PPA across variations of brainstem-pupil lags. Squares: unique PPA, that is PPA minimized over brainstem-pupil lags of the brainstem region of non-interest (y-axis). **g,** Mean ± s.e.m. unique/shared PPA in LC-NA and DR-5HT obtained after/without regressing-out the contribution of the other brainstem region to pupil size. Horizontal colored bars: significant pupil-predictive activity (cluster-based permutation tests, *P* < 0.05, *). Stacked plot: proportion of shared/unique PPA. Grey areas: zoom-in lags in **f**.

While these stratified analyses were performed at a single brainstem-pupil lag (maximizing brainstem-pupil correlation; individually identified in each animal), there may be a more complex interaction between brainstem systems at different lags. Thus, we investigated all possible combinations of brainstem-pupil lags. Specifically, we assessed whether brainstem systems independently, thus additively, modulated pupil size or whether pupil fluctuations reflected the multiplicative interaction between brainstem structures (e.g., as in a gating scenario). Additive and multiplicative combination captured similar patterns of pupil dynamics, but the additive combination resulted in larger pupil-predictive activity over all brainstem-pupil lags considered (Fig. 4c). This analysis is consistent with the notion that each brainstem system uniquely contributes to pupil dynamics.

Finally, we aimed to directly probe the strength of pupil-predictive activity in one system after removing the contribution of the other system. To do so, we obtained residual pupil dynamics, after removing (via linear regression) the contribution of one brainstem system, and then regressed the activity of the other system on these pupil residuals (Fig. 4d). For each brainstem-pupil lag considered, we report the minimum pupil-predictive activity identified from regression models ran across all past lags (Fig. 4e). This approach used more distant lags for DR-5HT than for LC-NA, in line with the observation that LC-NA drives DR-5HT (Fig. 2) and that pupil-predictive activity peaks first in LC-NA (Fig. 3). In addition, simulations indicated that this approach could reveal unique contribution to pupil changes in presence of shared covariation between brainstem regions (Supplementary Fig. 11). The approach identified unique contributions to pupil size from both LC-NA and DR-5HT (2.3 and 5.9%; Fig. 4f) over and above the impact of their correlated activity. Furthermore, the time course of unique contribution peaked earlier in LC-NA than DR-5HT (1 vs. 0.6s before), confirming previous results (Fig. 3). Using more distant time lags (20s instead of 10s) at which both brainstem regions do not even residually covary with pupil size anymore yielded identical conclusions, confirming the unique contribution of both LC-NA and DR-5HT to pupil dynamics (Supplementary Fig. 12). In sum, DR-5HT regulation of pupil size is not entirely mediated by LC-NA (as in *H*_2_): DR-5HT uniquely predicts pupil changes left unexplained by LC-NA.

### Activation of either LC-NA or DR-5HT dilate the pupil

Our results so far are in line with an LC-NA-independent contribution of DR-5HT to pupil dynamics. However, the results could also be explained by a third pupil-controlling region modulating DR-5HT activity. To assess and compare the contribution of LC-NA and DR-5HT to pupil control more directly, we optogenetically activated either LC-NA or DR-5HT in different mice (*DBH*- or *SERT*-cre respectively) using the same activation protocol (ChrimsonR; Fig. 5a).

**Fig. 5.**
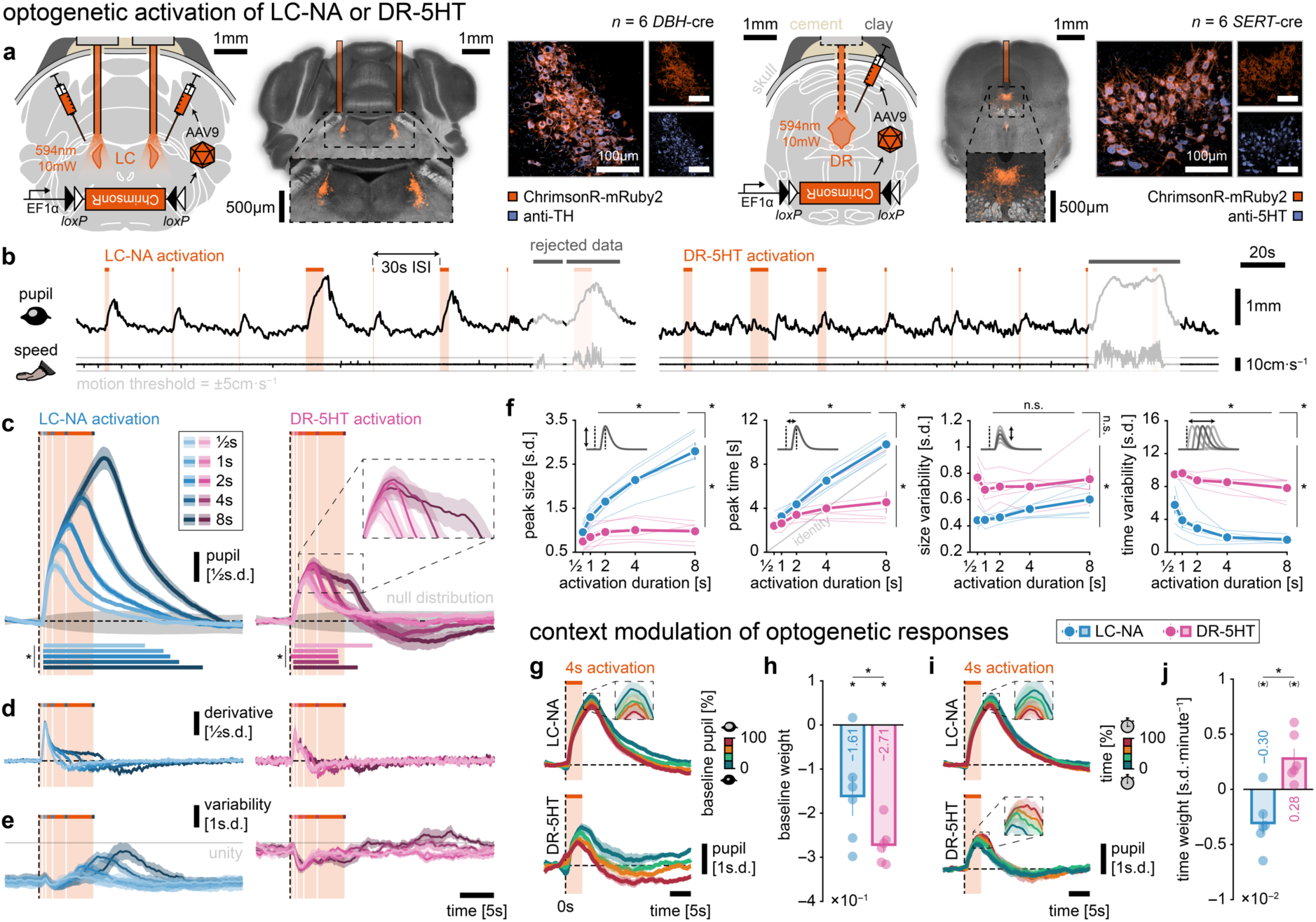
Activation of either LC-NA or DR-5HT dilate the pupil. **a,** The LC/DRN of *DBH*-/*SERT*-cre mice (*n* = 2 groups × 6 mice) were injected with the cre-dependent optogenetic excitatory tool ChrimsonR whose expression is restricted to NA/5HT cells respectively. The opsin was activated with orange light through implanted fiber optic implants. Expression of ChrimsonR in LC-NA/DR-5HT was confirmed with histology and immunostaining against TH/5HT. **b,** Snapshot of pupil size and speed in response to LC-NA or DR-5HT optogenetic activation (shaded orange areas). Trials entailing locomotion (speed > 5cm·s^−1^) were rejected. **c,** Mean ± s.e.m. pupil response to optogenetic activation of LC-NA or DR-5HT. Horizontal colored bars: pupil significantly larger than zero (cluster-based permutation tests, *P* < 0.05, *). Grey areas: bootstrapped null distribution of evoked responses (ɑ = 0.05, 10,000 permutations per session). **d,** Mean ± s.e.m. of the derivative of pupil response to optogenetic activation of LC-NA or DR-5HT. **e,** Mean ± s.e.m. trial-to-trial variability in pupil responses to optogenetic activation of LC-NA or DR-5HT. **f,** Mean ± s.e.m. peak response, time of peak, peak size variability and peak time variability. Significant modulation by activation duration (top), activated brainstem region (right) and interaction between them (corner; ANOVA, *P* < 0.05, *; n.s.: non-significant). **g,** Mean ± s.e.m. pupil response to 4s optogenetic activation of LC-NA or DR-5HT (as in **c**) split in quartiles of baseline pupil size (at *t* = 0s w.r.t. activation onset) after regressing-out the effect of time (see **i-j**). **h,** Regression weight of the baseline effect on pupil response (difference between peak and baseline). Coefficient significantly different from zero (Wilcoxon signed rank test, *P* < 0.05, *) and significantly different between LC-NA and DR-5HT (Wilcoxon rank sum test, *P* < 0.05, *) **i,** Same as **g** after a split in quartiles of activation time within a session after regressing-out the effect of baseline (see **g-h**). **j,** Same as **h** for activation time. Coefficient significantly different from zero (Wilcoxon signed rank test, *P* < 0.05 one-tailed, (*)) and significantly different between LC-NA and DR-5HT (Wilcoxon rank sum test, *P* < 0.05, *).

As expected^17–19^, LC-NA activation produced pupil dilations that scaled with activation duration (Fig. 5b-f). We also observed short-latency pupil dilations upon optogenetic activation of DR-5HT. Light delivery in non-opsin expressing animals did not alter pupil size, ruling out the possibility that luminance-induced pupil constriction would have masked part of the optogenetically evoked responses in opsin-expressing animals (Supplementary Fig. 13). The DR-5HT-induced pupil dilations were smaller and scaled less strongly with the duration of the activation than those induced by LC-NA (in the ranges we tested). This difference was not a simple light-dose effect due to the bilateral activation of LC: pupil dilations induced by unilateral activation of LC-NA were still larger than those observed after DR-5HT activation (Supplementary Fig. 13). Notably, we also observed that DR-5HT silencing induced pupil constriction, suggesting DR-5HT regulates pupil dynamics in a bidirectional manner (Supplementary Fig. 14).

We did not observe differences in response onset between LC-NA vs. DR-5HT systems, which could have indicated a hierarchical organization of these brainstem structures (Supplementary Fig. 15). Optogenetic activation of LC-NA or DR-5HT in our head-fixed mice neither initiated nor interrupted locomotion (Supplementary Fig. 16), thereby confirming that pupil responses are not the mere consequence of locomotion. Related, optogenetically induced pupil dilations were similar under isoflurane anesthesia, ruling out motion-/fear-/surprise-induced dilations or as the consequence of the subjective perception of brain manipulations^70,71^ (Supplementary Fig. 13).

A common assumption is that the more directly (i.e., with less intermediate relays) a stimulated brain structure controls the measured output, the smaller should be response variability. We observed that, although both LC-NA and DR-5HT activation quenched trial-to-trial pupil variability at activation onset, variability in the optogenetic responses was consistently larger for DR-5HT than for LC-NA (Fig. 5b-f). By contrast, pupil variability for LC-NA was confined to pupil relaxation when optogenetic activation was lifted and LC-NA recovered physiological variations.

We then aimed to isolate deterministic variables causing variability in optogenetic responses. First, we found a negative effect of baseline pupil on optogenetically evoked pupil response, with larger dilations happening on more constricted pupils. While this effect was observed for both brainstem structures, it was more pronounced in DR-5HT (Fig. 5g-h and Supplementary Fig. 17). Second, given the role of long-term effects and tonic modulation by LC-NA and DR-5HT^19,72,73^, we investigated the effect of time on optogenetic responses and observed a dissociation between brainstem structures. While pupil responses to optogenetic activation decreased over time in LC-NA, in line with a recent report^74^, they tended to increase over time in DR-5HT (Fig. 5i-j and Supplementary Fig. 17). In sum, while activation of both structures produces pupil dilation, the LC-NA control of the pupil seems stronger and less variable than the DR-5HT control. The latter might involve a more complex poly-synaptic pathway than the former.

### DR-5HT-induced pupil changes are only partially explained by LC-NA

Our previous experiments demonstrated a close relationship between DR-5HT activity and pupil size, but they did not test whether the DR-5HT control of pupil size depends (completely) on the concomitant activation of LC-NA (*H*_2_ vs. *H*_3_). We performed several analyses and experiments to arbitrate between these two hypotheses. In a first approach, we recorded LC-NA activity (jGCaMP7b) during DR-5HT optogenetic manipulations (ChrimsonR; Fig. 6a), while ensuring minimum cross-talk between recording and activation (Supplementary Fig. 18). DR-5HT activation produced a transient increase in LC-NA activity (Fig. 6b-c), the amplitude of which was largely independent of the duration of DR-5HT activation (Fig. 6d), a pattern reminiscent of DR-5HT-induced pupil changes (compare with Fig. 5). Moreover, the amount of LC-NA activity and pupil dilation induced by DR-5HT activation was of the same order of magnitude (Fig. 6e), and smaller than the effect of LC-NA activation on pupil (Fig. 6f). Thus, DR-5HT does drive LC-NA activity to some extent, which could explain how DR-5HT exerts its control on pupil dynamics.

**Fig. 6.**
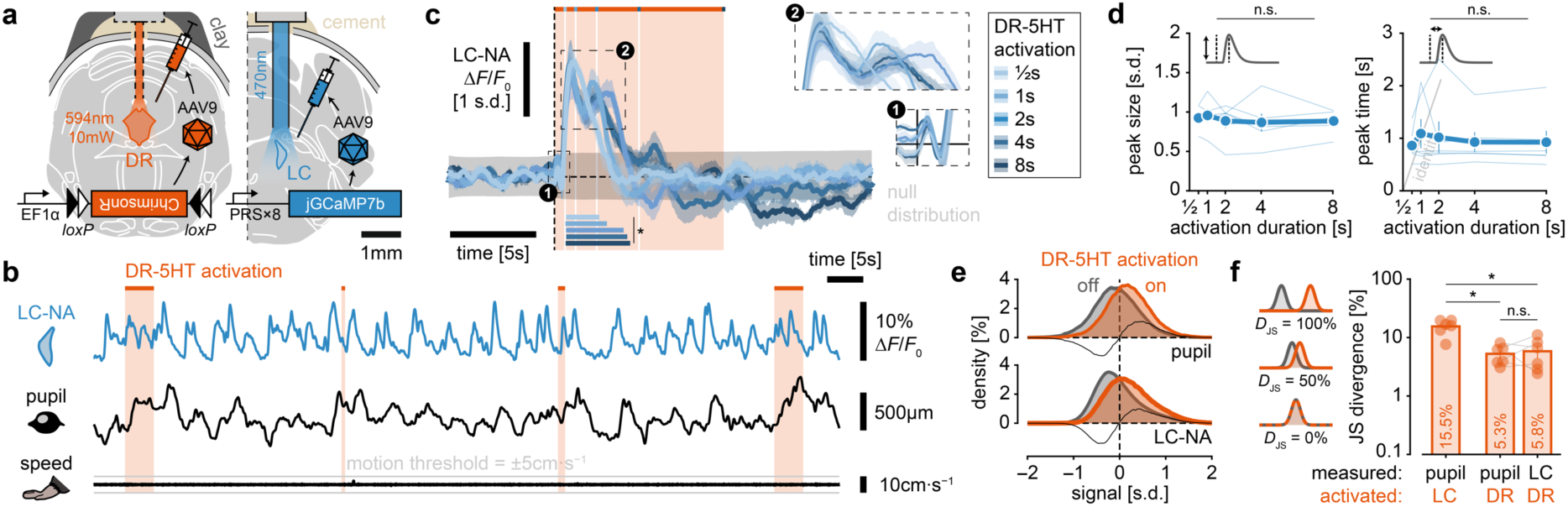
Activation of DR-5HT increases LC-NA activity. **a,** An optogenetic excitatory tool (ChrimsonR) and a calcium indicator (jGCaMP7b) were expressed selectively in LC-NA and DR-5HT respectively (*n* = 6 *SERT*-cre mice). **b,** Snapshot of LC-NA activity and speed in response to DR-5HT optogenetic activation. **c,** Mean ± s.e.m. LC-NA response to DR-5HT optogenetic activation. Horizontal colored bars: LC-NA activity significant different from zero (cluster-based permutation tests). Grey areas: bootstrapped null distribution of evoked responses (ɑ = 0.05, 10,000 permutations per session). **d,** Mean ± s.e.m. peak response and time of peak. No significant modulation of LC-NA activity by activation duration (ANOVA, *P* < 0.05, n.s.: non-significant). **e,** Mean ± s.e.m. distribution of pupil size (top) and LC-NA activity (bottom) in response to DR-5HT activation (on) vs. remaining samples (off). ‘On’ samples are defined as those belonging to significant temporal clusters (cluster-based permutation tests, *P* < 0.05; i.e., horizontal, colored bars in **c** and Fig. 5C). Thin line: difference between the two distributions. **f,** Mean ± s.e.m. Jensen-Shannon divergence (*D*_JS_) quantifying the overlap of ‘on’ and ‘off’ distributions for each pair of activated region/measured signal. Significant/non-significant differences (Wilcoxon signed rank/rank sum test, *P* < 0.05, *; n.s.: non-significant).

In a next approach, we tested whether the pupil responses evoked by DR-5HT activation could be explained by the concomitantly measured increase in LC-NA activity. We convolved spontaneous LC-NA activity with a pupil impulse response function (IRF; see Methods) and assess to which extent this LC-based model explained the pupil changes induced by DR-5HT activation (Fig. 7a). We first estimated the IRF parameters on sessions without any optogenetic stimulation; in other words, based on the relationship between spontaneous LC-NA activity and pupil dynamics (Fig. 7b-c). The cost of cross-session generalization of the thus-fitted IRF was remarkably low (< 1%; Fig. 7d; for details see Methods and Supplementary Fig. 19). In sum, our modelling approach captured meaningful and reproducible patterns of LC-pupil coupling.

**Fig. 7.**
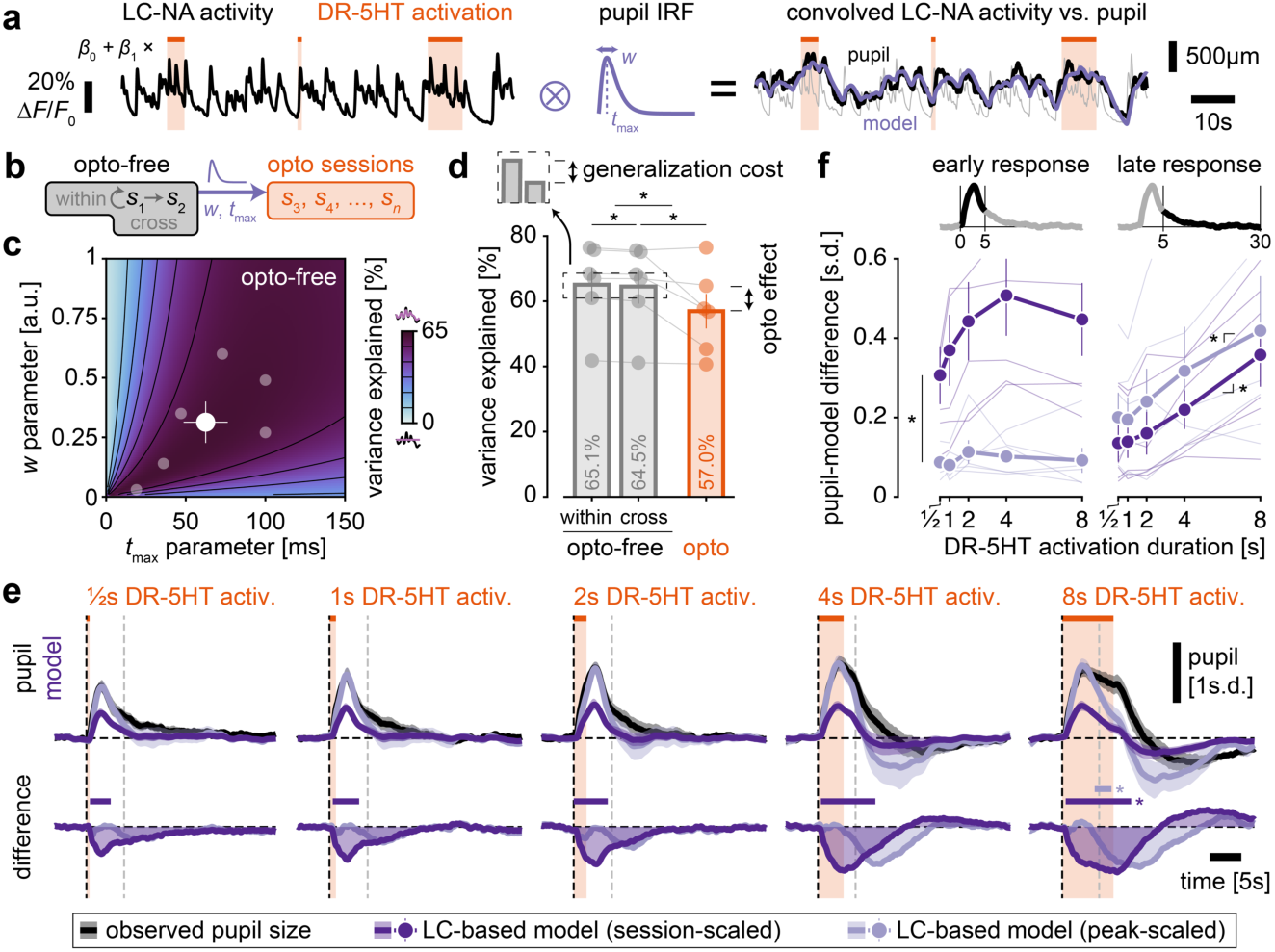
DR-5HT-induced pupil changes are only partially explained by LC-NA activity. **a,** LC-NA activity recorded during DR-5HT activation is convolved with the canonical pupil IRF and compared against true pupil size after scaling over the entire session. **b,** Parameters of the IRF (*w* and *t*_max_) are fitted in two opto-free sessions and applied to sessions with optogenetic manipulations. **c,** Mean proportion of variance explained and mean ± s.e.m. (filled, white dot) best-fitted IRF parameters (*w* and *t*_max_) in the two opto-free sessions. Semi-transparent dots: individual mice. **d,** Mean ± s.e.m. proportion of variance explained across different pairs of training vs. testing sets. Variance explained is either measured on the same opto-free session used for parameter fitting (within opto-free), on a different opto-free session than the one used for parameter fitting (cross opto-free) or measured in opto sessions using fitted parameters from both opto-free sessions. Semi-transparent dots: individual mice. Significant differences (Wilcoxon signed rank test, *P* < 0.05, *). **e,** Mean ± s.e.m. pupil response (as in Fig. 5) to DR-5HT activation and predictions of the LC-based model, either scaled to the level of the whole session or scaled to the peak of pupil evoked response separately for each activation duration. Second row: difference between model and the observed pupil. Horizontal colored bars: significant difference between pupil and the instances of the LC-based model (cluster-based permutation tests, *P* < 0.05, *). Vertical dashed grey line: limit between early and late responses (see **f**). **f,** Mean ± s.e.m. absolute error of the two instances of the LC-based model in two time-windows: early (0 to 5s; left plot) and late (5 to 30s, right plot). Thin lines: individual mice. Significant difference between instances of the LC-based model (left plot); significant modulation of model error by activation durations (right plot; Wilcoxon signed rank test, *P* < 0.05, *).

Applying this LC-based model to sessions with optogenetic activation of DR-5HT yielded a drop in variance explained (∼8% vs. < 1% for cross opto-free sessions). We also inspected predictions of the LC-based model specifically in time windows during the optogenetic activation of DR-5HT (Fig. 7e). The LC-based model did predict pupil dilations upon DR-5HT activation; however, the predictions were smaller in amplitude than the observed pupil (Fig. 7f). Moreover, the LC-based model less well predicted pupil changes after long DR-5HT activation duration, possibly indicative of the recruitment of pathways over and above the LC-to-pupil pathway.

The LC-based model might systemically underestimate pupil dilations (see Supplementary Fig. 19), thereby explaining the prediction of smaller-than-expected pupil responses also to optogenetic activation. We addressed this concern by rescaling the LC-based model’s predictions to the peak of opto-locked pupil responses and quantifying the resulting model’s error. Indeed, this did reduce the error in the prediction of the early part of the pupil response (around the peak), but the peak-scaled LC-based model still failed to explain the full dynamics of pupil responses to DR-5HT activation. Again, the late component of DR-5HT activation-induced pupil responses was not captured by the model (Fig. 7e-f). Together our analyses indicate that while part of the pupil responses induced by DR-5HT-activation were mediated by LC-NA, a component of induced pupil responses were unexplained by LC-NA dynamics. These observations are consistent with *H*_3_.

### LC-NA silencing dampens but does not abolish DR-5HT-incuded pupil dilations

As a final, critical test of *H*_3_, we monitored pupil dynamics in response to the optogenetic activation of DR-5HT during concomitant, optogenetic silencing of LC-NA. To this end, we used spectrally distinct optogenetic tools in the same mice (LC: PRS×8-Jaws, DR: cre-dependent ChR2(HR134R) in *SERT*-cre mice; Fig. 8a). The red-shifted chloride pump Jaws was used to maximize the extent of LC-NA silencing by strong and constant membrane hyperpolarization^75^ thereby suppressing action potentials^76^. Even though we used spectrally distinct optogenetic actuators, red-shifted opsins show residual sensitivity to blue light. We therefore aimed to minimize the extent of cross-activation in our experiment (Supplementary Fig. 20).

**Fig. 8.**
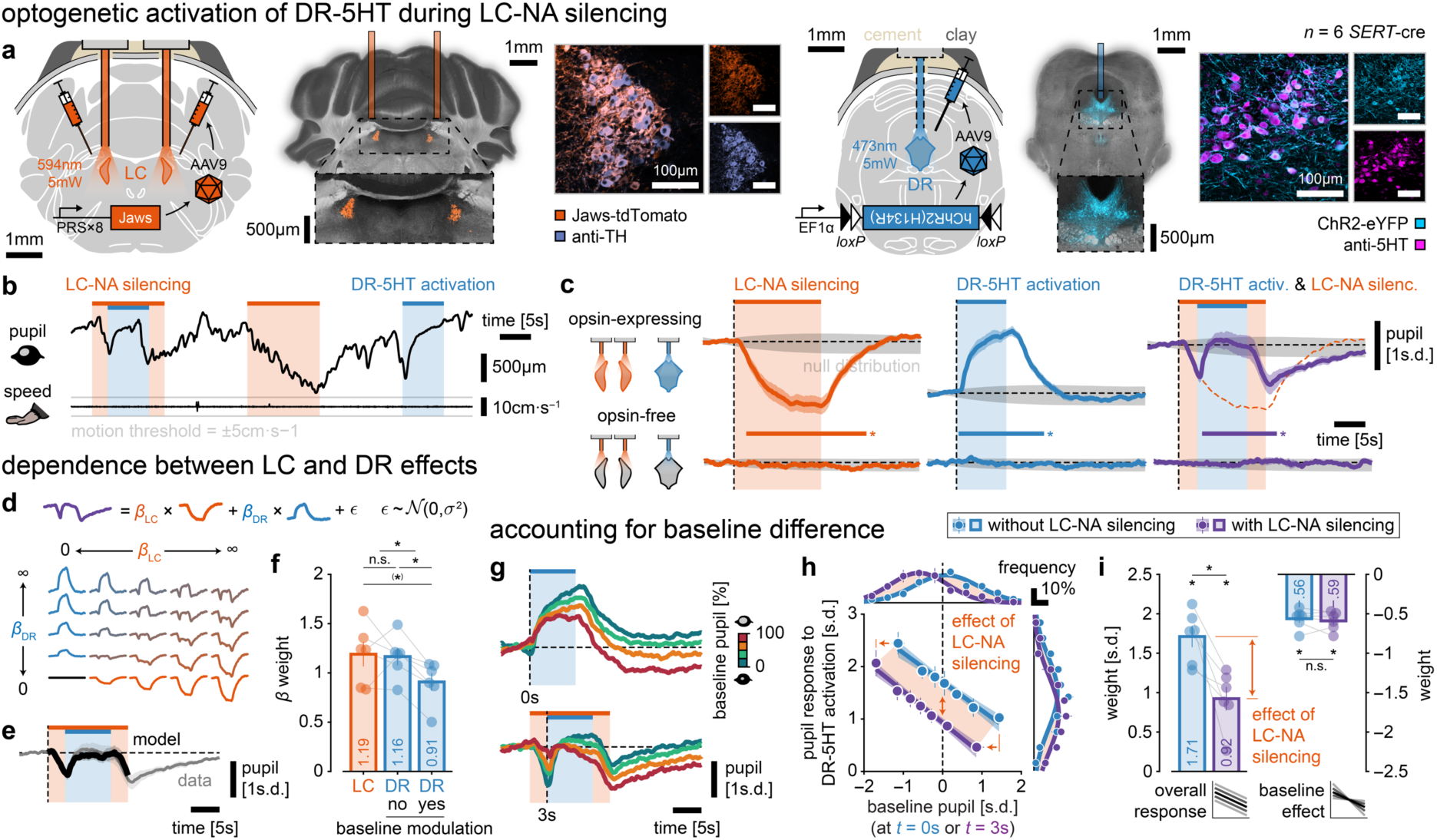
LC-NA silencing dampens but does not abolish DR-5HT-incuded pupil dilations. **a,** In *SERT*-cre mice (*n* = 6 mice), the LC was injected with the silencing tools PRS×8-Jaws and the DR with the cre-dependent optogenetic excitatory tool ChR2. Opsins were activated with orange and blue light respectively. Expression LC-NA/DR-5HT was confirmed with histology and immunostaining against TH/5HT. **b,** Snapshot of pupil size and speed in response to LC-NA optogenetic silencing and/or DR-5HT optogenetic activation. **c,** Mean ± s.e.m. pupil response evoked by LC-NA silencing, DR-5HT activation and DR-5HT activation during LC-NA silencing for opsin expressing animals (top row) and opsin-free animals (bottom row). Horizontal colored bars: response significantly larger than zero (left and middle time courses) or significantly different from LC-NA silencing (right time courses; cluster-based permutation tests, *P* < 0.05, *). Grey area: bootstrapped null distribution of evoked responses (ɑ = 0.05, 10,000 permutations per session). **d,** The pupil response to joint LC-NA silencing and DR-5HT activation is described as the linear, weighted sum to the pupil response to each optogenetic manipulation in isolation (positive weights only). **e,** Mean ± s.e.m. model prediction and observed pupil resulting from joint LC-NA silencing and DR-5HT activation. Only time course during light delivery is fitted. **f,** Mean ± s.e.m. weights for the contribution of LC-NA and DR-5HT (with and without baseline modulation, see **h-i**) in explaining pupil response to joint LC-NA silencing and DR-5HT activation. Significant differences in weights (Wilcoxon signed rank test, *P* < 0.05 two-tailed, *, one-tailed, (*); n.s.: non-significant). Semi-transparent dots: individual mice. **g,** Mean ± s.e.m. pupil response to optogenetic activation of DR-5HT with/without LC-NA silencing (as in **c**) split in quartiles of baseline pupil size (at *t* = 0s w.r.t. DR-5HT activation onset). **h,** Mean ± s.e.m. pupil response (difference between peak and baseline) to DR-5HT activation as a function of mean ± s.e.m. baseline pupil size with/without LC-NA silencing, and linear fit ± 95% confidence interval. LC-NA silencing restricts the range of baseline pupil size to more constricted pupils and dampens the overall pupil response. **i,** Mean ± s.e.m. weights of overall pupil response and baseline effect. Semi-transparent dots: individual mice. Significant differences in weights against zero and between conditions with/without LC-NA silencing (Wilcoxon signed rank/rank sum test, *P* < 0.05; n.s.: non-significant).

In accordance with previous studies^17,18^, silencing LC-NA caused pupil constrictions (Fig. 8b-c), confirming the viability of our approach. Optogenetic activation of DR-5HT in the absence of LC-NA silencing produced pupil dilations, replicating results from Fig. 5. If pupil responses induced by DR-5HT activation were mediated by LC-NA, silencing LC-NA should abolish these responses. In contrast, DR-5HT activation during LC-NA silencing still produced pupil dilations on top of LC-NA-induced pupil constrictions (Fig. 8b-c), in line with *H*_3_. Given that blue light is here delivered only during periods in which LC-NA is already silenced, cross-activation cannot account for the above result.

To assess the linear superposition of pupil constrictions induced by LC-silencing and pupil dilations induced by DR-activation, we then modeled the pupil response as a linear, weighted sum of pupil responses to individual optogenetic manipulations (LC silencing or DR activation, respectively; Fig. 8d-f; see Methods). The fitted weights were statistically undisguisable between LC and DR (1.19 vs. 1.16; Fig. 8g), in line with a linear superposition of the LC-NA and DR-5HT effects. Interestingly, this equal balance also results in the pupil returning to baseline levels after the pupil responses to both LC-NA silencing and DR-5HT activation have stabilized. While this effect could be explained by a perfect counterbalance of LC-driven silencing and DR-driven activation of the pupil apparatus system, the role of stimulation parameters (duration and intensity) and optogenetic tools (kinetics, potency, etc.) are likely central to this observation. Thus, we refrain from interpreting the absolute size of dilations under joint LC/DR manipulations.

The previous analysis however does not consider the inverse dependence between optogenetically induced pupil responses and baseline pupil size that we observed upon DR activation alone (Fig. 8g-i and Fig. 5). Thus, DR-induced dilations should be larger under LC-NA silencing (due to the associated pupil constriction) compared to sole DR-5HT activation. Relating baseline pupil (at DR-5HT activation onset) to the pupil response showed a negative linear effect that was of similar size with and without LC-NA silencing (0.56 vs. 0.59; Fig. 8i). Accounting for baseline pupil revealed that DR-5HT-induced pupil changes were dampened when LC-NA was silenced (1.71 vs. 0.92 s.d.). Note that incomplete LC-NA silencing would produce a pattern of results opposite to what we observed: larger pupil response to DR-5HT activation under poor LC-NA silencing when the pupils are less constricted. The fact that pupil responses to DR-5HT activation are dampened but still persisted during LC-NA silencing thus indicates that DR-5HT affects pupil dynamics both in an LC-NA-dependent and -independent manner, consistent with *H*_3_.

## Discussion

The ‘ascending reticular activating system’ was discovered decades ago, as a dense network of closely spaced, but neurochemically distinct nuclei^30–37^. The different neurochemical systems constituting this network have since then largely been studied in isolation because of limited experimental access to their interactions in awake animals. Elaborating their interactions, however, is critical for understanding the neuromodulatory control of brain states^15^. One line of work has studied the impact of the LC-NA and cholinergic systems on cortical states using pupil size as an established peripheral marker^4,6,22^. However, this research did not determine how these systems interact in regulating pupil and brain states. The understanding of the interplay between the DR-5HT and the LC-NA systems is of particular importance, because previous evidence points to distinct (partially even opposing) effects on central arousal and both systems are targeted by current treatments of mood disorders. Anatomy is compatible with several alternative modes of interactions because connections between LC and DR are dense and candidate anatomical pathways for pupil control by DR-5HT via the LC^30–37^ and bypassing the LC^77–79^ both exist, but each with as yet unknown functionality. Previous studies of the effects of DR-5HT stimulation on pupil size yielded complex, long-latency effects that are compatible with polysynaptic pathways and full mediation by the LC^28,29^. Simultaneous monitoring and independent manipulation of DR-5HT and LC-NA activity, concurrently with recordings of pupil size in awake animals was necessary to conclusively identify these functional interactions.

Here, we developed a multiplexed genetic targeting approach^46,47,80^ to assess this interaction in a comprehensive, neurochemically and anatomically precise manner in the awake mouse brain. Our approach enabled us to (i) study the natural interplay between the DR-5HT and the LC-NA systems, and (ii) determine their respective contributions to pupil-linked arousal through detailed analyses of their natural correlation structure as well as through precisely controlled manipulations. Critically, we identified joint and unique roles of both systems in regulating non-luminance mediated pupil dilations. Our study provides a first insight into the interplay of these two key neurochemical systems by revealing a tight, positive coupling between DR-5HT and LC-NA activity. With the combination of tools used here, we first confirmed the causal role of LC-NA in pupil dilation established by previous studies^16–18,21,22^. Specifically, we replicated the findings that (i) variations in LC activity shortly and uniquely predict subsequent changes in pupil size and (ii) activation or silencing of LC-NA dilates or constricts the pupil, respectively.

More importantly, we demonstrate both an LC-independent and an LC-dependent component DR-5HT-mediated pupil control. The following results indicate the existence of a DR-5HT pathway for pupil control, which bypasses the LC-NA system: (i) DR-5HT uniquely predicted patterns of spontaneous pupil fluctuations unexplained by LC-NA activity, (ii) pupil responses to DR-5HT activation were not entirely explained by changes in LC-NA activity, and (iii) DR-5HT-induced changes in pupil size persisted under LC-NA silencing. At the same time, our results also indicate that part of the DR-5HT pupil regulation is mediated indirectly, through the LC-NA: (i) LC-NA and DR-5HT spontaneous activities covaried and explained together a large fraction of pupil fluctuations, (ii) DR-5HT activation drove LC-NA activity, which explained part of DR-5HT-induced pupil dilations, and (iii) DR-5HT-induced pupil dilations were dampened during LC-NA silencing.

Taken together, our findings are consistent with a functional model (Fig. 1a, *H*_3_), in which LC-NA and DR-5HT systems interact closely to regulate pupil-linked arousal, but in which the DR-5HT system also makes a unique contribution. This model challenges widely held assumptions about the neuromodulatory control of pupil dynamics, and it is surprising in terms of the positive natural cooperation of the LC-NA and DR-5HT systems.

### Neural correlates versus neural drivers of pupil dilation

Pupil-related activity is observed in many cortical (such as the anterior cingulate cortex) and subcortical structures of the primate^16,21,23,60,81–84^ and rodent^53,85,86^ brain; a feature expected for a marker of brain state controlled by brainstem systems with diffuse ascending projections into the forebrain. Importantly, the number of structures that have so far been established to evoke pupil dilation at *short latency* upon direct stimulation is substantially smaller^7,10^. To date, the LC-NA system is considered the main central regulator of pupil fluctuations^16,17^. The close correspondence between LC-NA activity and pupil dynamics has often led researchers to use pupil size as a peripheral proxy of LC-NA activity^21,23^, thereby bypassing the difficulty of recording from deep brainstem structures such as the LC. Other central regulators of pupil dynamics include superior and inferior colliculi^16,87^, nucleus incertus^25^, basal forebrain^4,6,22^, and hypothalamus^27^. However, it has remained elusive whether these structures regulate pupil-linked arousal independently of LC-NA.

Here, we identified a unique contribution of DR-5HT to pupil dilation bypassing the LC-NA system. Thus, our study goes beyond first evidence for pupil reactivity to optogenetic activation of the DR-5HT in mice^28,29^. A direct, LC-NA-independent component can explain the short-latency, phasic pupil dilations that we observed in response to DR-5HT activation, which were more direct in nature and an order of magnitude faster than the modulations of pupil size under DR-5HT drive found previously (prolongation of a dilation existing even without DR-5HT stimulation emerging about 2s after stimulation onset)^28^. One practical implication of this insight is that any change in pupil size observed experimentally (or clinically) could be mediated by changes in only LC-activity, only DR-5HT activity, or a combination of both. The LC-NA-independent component may further account for pupil fluctuations that cannot be explained by changes in LC-NA activity^88,89^. Applying our approach to other neuromodulatory systems (e.g., dopamine) would allow the field to develop a comprehensive model of the relationship between brain states and pupil dilation, and to potentially identify temporally distinct components of pupil dynamics related to specific neuromodulatory systems.

### Broader implications

By revealing the interplay between the DR-5HT and LC-NA systems in awake animals, our findings have important implications for basic and translational neuroscience beyond the mechanistic interpretation of pupil dilation. The independent study of LC-NA and DR-5HT revealed striking parallels in the function of these two neuromodulatory systems. It is well established that the DR-5HT system plays important roles in the regulation of sleep-wake cycles^39–41^, neural plasticity^14^, and sensory processing^42–44,90^. It has been postulated that DR-5HT is involved in predictive processing and the monitoring of uncertainty or confidence even in the absence of reward or feedback^14^. Similar assumptions hold for LC-NA activity^1,2,45^, and pupil dilation^62,91–93^. Likewise, the DR-5HT and LC-NA systems are implicated in major neuropsychiatric disorders such as depression^1,14^. Correspondingly, both systems are the targets of widely used pharmacological treatments of such disorders, including selective serotonin or noradrenaline reuptake inhibitors.

Unravelling the functional interaction between these two monoaminergic systems is therefore critical for developing a mechanistic understanding of their role in both healthy cognition and neuropsychiatric disorders. Our study makes a first advance in this direction. The positive intrinsic and stimulation-induced coupling between LC and DR is at odds with studies using pharmacological application of 5HT in visual cortex, which causes a reduction of the gain of sensory neurons^44^, an effect opposite to the one observed for NA^1,94,95^. While this superficial discrepancy could be related to distinct effects of tonic vs. phasic 5HT action in the brain^14^, the close cooperation between these two major neuromodulatory systems in the control of waking state is a first, important finding and future work should now identify specific differences in their functions.

### Limitations and outlook

While our study dissected the functional interactions between LC, DR and pupil, it did not resolve the precise anatomical pathways mediating LC-dependent and LC-independent DR-5HT effects on pupil size. Pupil size depends on the activity balance between two cell groups, in the parasympathetic Edinger-Westphal nucleus (for constriction) and the sympathetic intermediolateral cell column of the spinal cord (for dilation)^96^, each of which receives inputs from many subcortical structures^7^, including LC and DR. The LC-dependent pupil regulation by DR-5HT may be indirect (via intermediate nuclei) or, as suggested by the short-latency effects, mediated by direct connections from the DR-5HT neurons to the LC^30–36^. Those connections could disinhibit the LC (i.e., by activating inhibitory 5HT_1a_ receptors in peri-LC GABAergic cells^17^) or directly drive LC activity (e.g., by releasing 5HT within LC^37^). By contrast, the LC-independent component could be implemented by a 5HT drive of the intermediolateral column of the spinal cord^7,10,20^ or indirectly via the hypothalamus^77–79^. Future work should dissect these pathways at a circuit level.

In addition, refined studies are needed in the future to evaluate to what extent different neuronal subpopulations within DR-5HT^97–104^ and LC-NA^105–110^ exert distinct pupil controls and project to different pupil-controlling regions. For example, activation of DR-5HT thalamic terminals does not affect the pupil^111^, suggesting pupil control requires synchronous activity of a large portion of the DR-5HT. Functional heterogeneity of 5HT-neuronal populations in the DR could possibly explain the slower dynamics we observed in DR-5HT compared to LC-NA, since our recordings only reflect the average activity of the entire population. As a consequence, the difference in dynamics between LC-NA and DR-5HT may partly underly the unexpected observation that pupil-predictive activity ramped earlier and was stronger in DR-5HT than in LC-NA, as well as the distinct coupling between LC vs. DR and pupil at high vs. low frequencies respectively. Recordings at cellular resolution from identified neuronal subpopulations will be needed in future studies to investigate the temporal dynamics in pupil-predictive activity of LC-NA vs. DR-5HT in more detail.

The existence of multiple subpopulations of DR-5HT may also reconcile the mydriasis-promoting role of DR-5HT we and others^28,29^ have reported, with other studies reporting an arousal-reducing role of DR-5HT^44,49,73,112,113^. Another possibility is that DR-5HT manipulations yield opposite effects when examined at different time scales^14,73^ and in different contexts^43^. One possible context-dependent effect concerns the regulation of locomotion: while locomotion is associated with strong pupil dilation preceded by brainstem activity, the optogenetic activation of LC-NA or DR-5HT did not trigger locomotion. One hypothesis is that arousal causes locomotion only in specific states, possibly regulated by downstream neuronal populations. Alternatively, locomotion could be triggered by higher-level structures that recruit other brain areas (e.g. basal ganglia) in parallel to LC and DR, such that LC and DR manipulations would be insufficient to trigger running. Future research could inspect the coupling between brainstem systems and the pupil in different behavioral states and in task-engaged animals.

Translating our findings to species other than mice requires caution. For example, activation of 5HT_1a_ receptor causes pupil dilation in mice^114^ but pupil constriction in marmosets^115^, a difference possibly explained by species diurnality^20^ or co-activation between DR-5HT and other neuromodulatory systems^116,117^. Arousal positively correlates with pupil size in mammals but not necessarily in all species (e.g., constriction during aroused wakefulness in birds^118^). Thus, our findings are likely to generalize from the mouse brain to the primate brain.

## Conclusion

In the current work, we have developed an approach for unraveling the interaction of two important brainstem systems, the DR-5HT and LC-NA systems, in controlling pupil-linked arousal states. Our approach uncovered a close, cooperative interaction of the phasic activity of both brainstem centers, both in the spontaneous activity and under external manipulation, as well as two short-latency components of pupil dilation controlled by DR-5HT: one via the LC and another one bypassing the LC. These insights can pave the way for the development of more refined theories of neuromodulatory control of brain states as well as more refined treatments of neuropsychiatric disorders.

## Data availability

The PRS×8-driven constructs engineered for this study will be made available to the community on *AddGene* upon publication: PRS×8-jGCaMP7b as plasmid #192594, PRS×8-jRGECO1a as plasmid #210399, and PRS×8-Jaws-tdTomato as plasmid #192593.

The datasets and analysis code are available upon request to the corresponding author.

## Acknowledgements

We thank Cynthia Rais for useful discussions and for demonstrating experimental procedures, Stefan Schillemeit for genotyping and admin support, Kathrin Sauter for plasmid cloning and admin support, Alexander Dieter for demonstrating experimental procedures, and Ingke Braren for virus production. We are also grateful to Zachary Mainen, Fanny Cazettes, Elisabete Augusto and members of the Mainen lab for useful discussions.

This work was supported by a *Humboldt postdoctoral individual fellowship*, by the *Bettencourt Schueller foundation* (both to M.M.) and by the *Deutsche Forschungsgemeinschaft* (*DFG*, *German Research Foundation*) SFB 936 – 178316478 - A7, Z3 (to T.H.D.) & B8 (to J.S.W.).

## Author contributions

M.M.: conceptualization, methodology, software, validation, formal analysis, investigation, data curation, writing — original draft, writing — review & editing, visualization, project administration, funding acquisition; T.H.D. & J.S.W.: conceptualization, writing — review & editing, funding acquisition.

## Additional information

**Supplemental information** is available.

**Correspondence** should be addressed to Maxime Maheu.

## Methods

### Subject details

#### Ethics

All experiments followed the 3R principles and were performed in compliance with German law according to the directive 2010/63/EU of the European Parliament on the protection of animals used for scientific purposes. The particular set of experiments presented in this study was approved by the Hamburg state authority for animal welfare under licenses #33/2019 and #005/2023.

#### Animals

Experiments were performed on 32 adult mice (24 females) aged 8 to 40 weeks (mean: 20 weeks) at the time of surgery. All mice were heterozygously expressing cre recombinase either in serotonin transporter (SERT) positive neurons^80^ (B6.129(Cg)-*Slc6a4^tm1(cre)Xz^*/J, referred to as *SERT*-cre; RRID: IMSR_JAX:014554) or dopamine beta hydroxylase (DBH) positive neurons^119^ (B6.Cg-*Dbh^tm3.2(cre)Pjen^*/J, referred to as *DBH*-cre; RRID: IMSR_JAX:033951). Mouse genotype was verified for each animal by polymerase chain reaction and gel electrophoresis.

#### Housing

All transgenic mice were bred from in-house colonies originally derived from two male mice obtained commercially (*The Jackson Laboratory*, USA). When mice reached adulthood, they were moved from the breeding facility to a dedicated animal room close to the experimental setups in order to minimize transfer-induced stress. Temperature (∼22°C) and humidity (∼40%) were maintained constant and a 12h/12h reversed light-dark cycle was used throughout the duration of experiments.

### Experimental procedures

#### Surgical procedures

Experiments required the local injection of reagents in multiple regions of the brainstem and the placement of fiber optic implants above the injected areas. All procedures were done in a single surgical procedure which typically lasted 1 to 2 hour(s). Throughout the surgery, the animal was placed under general anesthesia through inhalation of isoflurane (induction: 2% for at least 5 minutes, maintenance: ∼1.5%) mixed with air (RSA-A(H)-S; *RWD*, China). Analgesia was achieved by subcutaneous injection of Buprenorphine (0.1mg·kg^−1^ in NaCl). Anesthesia and analgesia were then confirmed by the abolition of the hind limb withdrawal reflex. The absence of reflex response was regularly checked throughout the surgery which, combined with breathing monitoring, served to fine tune isoflurane concentration. The animal was placed on a heating pad maintaining body temperature (at 37°C) through a closed-loop system combining an anal temperature probe and a heating device (69027; *RWD*, China). In addition, temperature was also occasionally checked with an infrared thermometer (IR500-12S; *Voltcraft*, UK). Upon confirmed anesthesia and analgesia, the animal was attached to a digital stereotactic surgical frame (100μm precision, 51725D; *Stoelting*, USA) using an auxiliary ear bar (EB-6; *Narishige*, Japan) to minimize damage to the animal’s ear tube and eardrum. The scalp was shaved using a trimmer (MT5531; *Grundig*, Germany) before being disinfected with iodine solution (Betaisodona; *Mundipharma*, Germany). Both eyes were then covered with eye ointment (Vidisic; *Bausch & Lomb*, Germany) to avoid drying. An antero-posterior (AP) skin incision was then made (∼1.5cm long), and the skull was cleaned from any remaining tissue. Anatomical landmarks bregma and lambda (taken as −4.1mm posterior to bregma) were aligned along the dorso-ventral (DV) and medio-lateral (ML) axes. Additional lateral landmarks (±2mm lateral to bregma) were further aligned along the DV axis. Craniotomies were made (∼500μm diameter) using a rotary micromotor (∼6300rpm; K.1070; *Foredom*, USA) equipped with a steel drill (310 104 001 001 021; *Meisinger*, Germany): one above each LC and one above the DR. A glass micropipette was pulled from a borosilicate glass capillary (B100-20-10; *Sutter Instruments*, USA) using a micropipette puller (PC-100; *Narishige*, Japan). The micropipette was then filled with a suspension of recombinant adeno-associated viral (AAV) particles diluted in phosphate-buffered saline (PBS) solution. The filled micropipette was slowly lowered into the LC (±1mm ML, −5.4mm AP, −3.6mm DV to bregma) and into the DR with a 32° degree posterior angle^50^ (after trigonometric transform: −6.5/−6.5/−7.0mm AP, and −4.0/−3.6/−3.9mm DV to bregma). Note that LC injection coordinates were slightly more lateral to LC location to minimize the risk of injecting into the nearby 4^th^ ventricle. Moreover, injections were made at 3 different sites within the DR to cover its entire volume and to mitigate the risk of injecting into the nearby 4^th^ ventricle. The viral suspension was slowly expelled (at a rate of ∼200nL·min^−1^) using a custom-made manual air pressure system until the desired volume was reached (∼300nL per LC; ∼300nL per injection site in DR). The micropipette was kept in place for at least 1 min before it was slowly retracted. Once all injections were completed, optic fiber stubs were implanted straight above the LC (calcium recordings: ±0.9mm ML, −5.4mm AP, −3.5mm DV; optogenetic manipulations: ±1.0mm ML, −5.4mm AP, −3.4mm DV) and above the DR using a 32° posterior angle^50^ (after trigonometric transform: −6.7mm AP, −3.6mm DV for calcium recordings and −3.5mm DV for optogenetic manipulations). For optogenetic manipulations, 200μm core diameter fiber optic implants were used (MFC_200/245-0.37_3.7mm_ZF1.25g_FLT; *Doric Lenses*, Canada), while 400μm ones were used for calcium recordings (MFC_400/430-0.37_3.7mm_ZF1.25g_FLT; *Doric Lenses*, Canada), both with 0.37 numerical aperture (NA). Craniotomies were sealed and fiber optic implants were maintained in place using tissue adhesive (1469SB, *Vetbond*; *3M*, USA) which was also used to attach the skin onto the skull. The skull and the base of the fiber optic implants were then covered with opaque dental cement (*Super-Bond* universal kit; *Sun Medical*, Japan). A head bar (200-200 500 2110; *Luigs & Neumann*, Germany) was finally implanted anterior to the fiber optic implants and further immersed in dental cement. An intraperitoneal injection of Carprofen (4 mg·kg^−1^, diluted in NaCl) was performed as a post-surgery analgesic and anti-inflammatory treatment. The mouse was finally removed from the stereotaxic frame and allowed to recover alone in a dedicated cage partially placed on top of a heating pad. The animal was carefully monitored throughout the wakening stage and was placed back into its home cage upon recovery (as indicated by extensive locomotion). Meloxicam (0.5mg·mL^−1^; *Metacam*; *Boehringer Ingelheim*, Germany) was mixed into softened food for the three days following surgery.

#### Viral constructs

Both LC-NA and DR-5HT were targeted in the same mice using a combined strategy: on the one hand, expression was restricted to 5HT cells (sensitivity ≈ 1^80^; specificity = 0.94^49^ and 0.99^80^) using the cre-recombinase technique with floxed constructs in *SERT*-cre animals^80^; on the other hand, expression was restricted to NA cells (sensitivity = 0.78; specificity = 0.62^47^) thanks to the synthetic promoter PRS×8 derived from the human DBH promoter^46^. Note that alternative strategies could have also been used: a synthetic promoter for targeting 5HT cells (e.g., TPH2) combined with a mouse line for NA cells (e.g., *DBH*-cre); or crossing of two mouse lines each specialized for one cell type (e.g., *DBH*-cre with *SERT*-cre). These alternative strategies however have constraints: poorly characterized transduction efficiency or risk of cross-transduction across regions given the anatomical proximity of LC and DR. As a consequence, we opted for a combined, promoter/mouse-line strategy (PRS×8/*SERT*-cre). For DR injections, transgenes were packaged in AAV2/1 and AAV2/9 capsids, both of which have high tropism for 5HT cell populations^120^. For LC, we used exclusively AAV2/9 capsids which also provide efficient transduction of NA cell populations^47^. For calcium recordings, we used one of two approaches: injecting either pAAV2/1-Syn-Flex-jGCaMP7b (ITR-derived titer = 1.40 × 10^13^ vg·mL^−1^; *AddGene* viral prep #104493-AAV1; a gift from Douglas Kim & GENIE Project^121^) in DR and pAAV2/9n-PRS×8-jRGECO1a (ITR-derived titer = 8.5 × 10^13^ vg·mL^−1^) in LC, or pAAV2/1-Syn-Flex-NES-jRGECO1a (ITR-derived titer = 2.6 × 10^13^ vg·mL^−1^; *AddGene* viral prep #100853-AAV1; a gift from Douglas Kim & GENIE Project^122^) in DR and pAAV2/9-PRS×8-jGCaMP7b (ITR-derived titer = 1.0 × 10^15^ vg·mL^−1^) in LC. For LC recordings, injection was done bilaterally and recording was done unilaterally from the LC exhibiting the strongest calcium transients on the first acquisition day, and maintained throughout all acquisition sessions. Because blue light promotes arousal in mice (through melatonin release suppression)^123^, we opted for red-shifted opsins whenever possible. For LC and DR single-region optogenetic activations, we used pAAV2/9n-EF1α-DIO-ChrimsonR-mRuby2-Kv2.1 (ITR-derived titer = 9.0 × 10^12^ vg·mL^−1^; *AddGene* viral prep #124603-AAV9; a gift from Christopher Harvey^124^). For joint DR activation and LC recording, the PRS×8-jGCaMP7b and the DIO-ChrimsonR constructs listed above were used. For joint LC-DR optogenetic manipulations, we used pAAV2/9n-EF1α-DIO-hChR2(H134R)-eYFP (ITR-derived titer = 2.4 × 10^13^ vg·mL^−1^; *AddGene* viral prep #20298-AAV9; a gift from Karl Deisseroth^125,126^) in DR and pAAV2/9-PRS×8-Jaws-KGC-tdTomato (ITR-derived titer = 2.0 × 10^15^ vg·mL^−1^; modified from *AddGene* plasmid #153538; a gift from Edward Boyden^76^) in LC. For both LC activation (with ChrimsonR) and LC silencing (with Jaws), injection and light delivery were done bilaterally.

#### Experimental setup

The animals recovered for at least two weeks after the surgery before they were habituated to the experimenter, the experimental setup, and to head fixation, in that order. After habituation, experiments were conducted between 3 to 8 weeks post-surgery. Mice were head fixed laterally from the right on a linear 7cm wide and 2m long treadmill (700-100 100 0010; *Luigs & Neumann*, Germany) which was placed within a noise-reducing custom-made box with acoustic foam walls. The setup was illuminated with stable, controlled ambient light (see pupil section below). Mice were allowed to voluntarily walk on the treadmill but were not rewarded to do so in order to minimize the occurrence of locomotion bouts.

#### Synchronization

Locomotion data, optogenetic stimulation trains, calcium recordings and eye video frames were all synchronized to a common timestamp system. For this purpose, input and output signal lines were plugged to the connector block (BNC-2110; *National Instruments*, USA) of a data acquisition board (PCIe-6323; *National Instruments*, USA) connected to a PCI-express slot of a computer (*Fujitsu Celsius* M770 with *Intel Xeon* W-2123 3.60GHz CPU, 16GB DDR4 2.666MHz RAM, SSD 512GB) operating *Windows* 10 Pro (version 10.0.19045 N/A Build 19045; *Microsoft*, USA). Acquisition was performed using custom-written *MATLAB* (*MathWorks*, USA) code at a sampling rate of 1800Hz.

#### Pupil recordings

The animal’s left eye was illuminated with an infrared 830nm light source and recorded with a monochrome, input-triggered camera (DMK 33UX249; *The Imaging Source*, Germany) equipped with a fixed-working-distance objective (TMN 1.0× 50mm; *The Imaging Source*, German) and a 780nm high-pass filter (FGL780; *Thorlabs*, Germany). Baseline pupil size was maintained at an intermediate level (neither too constricted not too dilated) by an ultraviolet LED oriented towards the right eye and whose brightness level was controlled by a micro-controller board (*Arduino Nano*; *Arduino*, Italy) through pulse width modulation. The acquisition of eye frames was triggered by a digital 30Hz pulse train generated by the data acquisition board to the camera.

#### Locomotion recordings

The animal’s position along the treadmill was monitored at a 20Hz sampling rate by combining digital signals from a rotary motion sensor (700-100 300 0902; *Luigs & Neumann*, Germany) and a lap reset optical sensor (700-100 300 0901; *Luigs & Neumann*, Germany).

#### Calcium recordings

Bulk activity of LC-NA and DR-5HT was recorded using fiber photometry of genetically encoded fluorescent calcium indicators (jGCaMP7b and jRGECO1a). Excitation and emission light travelled along the same optical path to and from the brainstem through the implanted fiber optic implant. Excitation light was provided by two multi-color LED light sources (pE-4000; *CoolLED*, USA) set to distinct excitation wavelengths (470nm for jGCaMP7b, 550nm for jRGECO1a). Excitation light power was set either to 50, 100 or 150μW at the fiber tip on an animal-per-animal basis depending on overall fluorescence strength, and stimulation parameters were held constant across acquisition sessions. Each light source was plugged to a fluorescence mini-cube (FMC5; *Doric Lenses*, Canada) housing dichroic mirrors with built-in bandpass excitation (400-480nm for jGCaMP7b, 555-570nm for jRGECO1a) and emission (555-570nm for jGCaMP7b, 580-680nm for jRGECO1a) filters. The output light path of the fluorescence mini-cube was connected to the fiber optic implant by a low auto-fluorescence patch-cord (MFP_400/440/LWMJ-0.37_1m_FC-ZF1.25f,_LAF; *Doric Lenses*, Canada) and a white 1.25mm zirconia mating sleeve (SLEEVE_ZR_1.25; *Doric Lenses*, Canada). Emitted fluorescence resulting from excitation of calcium indicators was detected with a photodetector system (gain = ×1; DFD_FOA_FC; *Doric Lenses*, Canada). Excitation lights and fluorescent read-outs were both controlled by a dedicated micro-controller board (pyBoard v1.1; *MicroPython*, USA) running a custom-modified version of the pyPhotometry software^127^ (with Python v.3.5) affording control of two separate light sources and two photodetectors. Because of the relatively close anatomical proximity between LC-NA and DR-5HT, and to avoid the risk of cross-talk, we sequentially recorded from each of these two regions using time-division multiplexing. Acquisition was done at a sampling rate of 90Hz for each of the two regions. In each acquisition sweep, background illumination was measured in a time window of 250μs immediately before excitation light was turned on for a duration of 750μs. Resulting fluorescence was measured in the last 250μs of the stimulation window and baseline subtracted. Only time periods where the animal was sitting quietly were kept for further analyses (see preprocessing below).

#### Optogenetic manipulations

Optogenetic stimulation light was generated by a laser combiner system (LightHUB; *Omicron*, Germany) housing 473nm (LuxX 473-100; *Omicron*, Germany) and 594nm diode lasers (Obis 594nm LS 100mW; *Coherent*, Germany). For singe-region optogenetic activation, the laser combiner system was connected to a multimode splitter branching patch-cord with two end ferrules (200μm core diameter, 0.22NA, 50:50 split; TT200SL1A, *Thorlabs*, Germany). For LC manipulations, each of the two ferrules was connected to each of the two cannulas implanted on top of each of the two LC. For DR manipulations, one of the two ferrules was connected to the single cannula implanted on top of the DR. Control experiments were performed to ensure differences between LC and DR did not arise from difference in the amount of light delivered (through 2 cannulas for LC vs. 1 for DR; see corresponding section below). For joint LC-DR optogenetic manipulations, the laser combiner system was connected to an asymmetric multimode splitter branching patch-cord (200μm core diameter, 0.22NA, 75:25 split; TM200R3S1A; *Thorlabs*, Germany). The high transmission end (75%) was connected to the input port of a multimode fiber-optic filter mount (FOFMS; *Thorlabs*, Germany) equipped with a 532nm high-pass filter (BLP01-532R-25; *Semrock*, USA). The output port of the filter mount was connected to a multimode splitter branching patch-cord with two end ferrules for bilateral LC silencing (200μm core diameter, 0.37NA, 50:50 split; SBP(2)_200/220/900/900-0.37_1m_SMA-2xZF1.25; *Doric Lenses*, Canada). The low transmission end (25%) was connected to the input port of a second multimode fiber-optic filter mount equipped with a 532nm low-pass filter (BSP01-532R-25; *Semrock*, USA). The output port of the filter mount was connected to a mono fiber-optic patch-cord with one end ferrule for DR activation (200μm core diameter, 0.37NA; MFP_200/220/900-0.37_1m_SMA-ZF1.25; *Doric Lenses*, Canada). In all cases, a white 1.25mm zirconia mating sleeve (SLEEVE_ZR_1.25; *Doric Lenses*, Canada) was used to couple the ferrules to the implanted fiber optic implants. Black modelling clay (29413F03; *Play-Doh*) was added around the mating sleeve and on top of the dental cement implant to restrain optogenetic light from inducing luminance-mediated pupil changes or being noticed by the animal. Control experiments were nevertheless performed to address this potential confound (see corresponding section below). Before each experiment, power at the tip of the fiber optic implant was calibrated to 5mW using an optical power meter (PM400; *Thorlabs*, Germany). Optogenetic stimuli consisted of 0.5, 1, 2, 4 or 8s continuous light pulses for single-region optogenetics, and 8 or 14s for joint LC-DR optogenetics. In all cases, a stimulation onset asynchrony of 30s was used. Stimulus trains were generated using custom-written *MATLAB* scripts (*MathWorks*, USA) controlling the digital acquisition board used for synchronization (see corresponding section above) connected to the digital inputs of the laser combiner system.

#### Optogenetic control experiments

The first control experiment addresses the possibility that optogenetically-induced pupil dilations were due to either (1) the subjective perception of brain manipulations, a mechanism previously identified in a wide range of manipulated brain regions and coined optoception^70^, or (2) to the subjective perception of thermal changes due to optogenetically-induced heat^71^. To this aim, we ran the LC vs. DR optogenetic activation protocol under isoflurane-anesthesia (∼1.5%) in opsin-expressing *DBH*- and *SERT*-cre animals (*n* = 2 × 6; same animals as those employed for the main experiment). The second control experiment addressed the possibility that larger pupil dilations evoked by LC optogenetic activation, in comparison to DR-induced ones, were caused by the bilateral nature of LC activation resulting in twice the amount of excitation light delivered in comparison to DR activation. To address this concern, we performed unilateral LC activation (either left or right) in opsin-expressing *DBH*-cre animals (*n* = 6; same animals as those employed for the main experiment). The third control experiment addressed the possibility that optogenetically induced pupil changes were caused, or at least affected, by luminance-mediated pupillary reflex in response to the laser light. To rule out this possibility, we performed the optogenetic light stimulation protocol (as described above) in *SERT*-cre animals expressing a cre-dependent fluorophore in DR and a PRS×8-promoted fluorophore in LC instead of the optogenetic actuators used in the main experiment (*n* = 6).

#### Perfusion

Upon completion of all recordings, brains were collected for histological verification of transgene expression. To this aim, mice were first deeply anesthetized with a lethal intra-peritoneal injection of ketamine and xylazine (180 and 24mg·kg^−1^ in NaCl, respectively). Upon abolition of the hind limb withdrawal reflex, the thorax was incised, the diaphragm opened and the thoracic cage removed. Mice were transcardially perfused, first with PBS (∼100mL) to rinse out the blood, then with paraformaldehyde (PFA; 4%) in PBS (∼100mL) to fix the tissue. The brain was then explanted and stored in 4% PFA at 4°C in a dark environment until further processing.

#### Immunohistochemistry

Brains were recovered, embedded in agarose (3%; Genaxxon *Bioscience*) and sliced coronally (70μm thickness) with a microtome (VT1000 S; *Leica*, Germany). Unspecific antibody binding site in slices were blocked in normal goat serum (NGS; 10%) and Triton X-100 (0.3%) in PBS for 2h at room temperature. Slices were then incubated with primary antibodies (with 10% NGS, 0.3% Triton X-100, in PBS) for 2 days at 4°C. NA-positive cells in LC-containing brain slices were labelled using an anti-tyrosine hydroxylase (TH) primary polyclonal antibody raised either in rabbit (1:500 dilution; AB152; *Merck Millipore*, USA) or in guinea pig (1:500 dilution; 213104; *Synaptic Systems*, Germany). 5HT-positive cells in DR-containing brain slices were labelled using either a primary polyclonal antibody from rabbit (1:250 dilution; 20080; *Immunostar*, USA) or goat (1:250 dilution; 20079; *Immunostar*, USA). jGCaMP7b and eYFP (fluorescent tag for ChR2) were amplified using a chicken anti-GFP primary polyclonal antibody (1:1000 dilution; A10262; *Invitrogen*, USA). jRGECO1a and tdTomato (fluorescent tag for Jaws) were amplified using a rabbit anti-DsRed polyclonal antibody (1:1000 dilution; 632496; *TaKaRa*, Japan). After incubation with primary antibodies, slices were rinsed 3 times for 5 minutes in PBS. Rinsed slices were then incubated for 1 day at 4°C in a carrier solution (2% NGS, 0.3% Triton X-100, in PBS) containing combination of the following secondary polyclonal antibodies (1:1000 dilution): (1) a goat anti-chicken Alexa Fluor 488nm (A11039; *Invitrogen*, USA), and/or (2) a donkey anti-rabbit Alexa Fluor 546nm (A10040; *Invitrogen*, USA), and/or (3) a goat anti-rabbit (A27040; *Invitrogen*, USA), or a donkey anti-goat (A21447; *Invitrogen*, USA), or a goat anti-guinea pig (A21450; *Invitrogen*, USA) Alexa Fluor 647nm. Transgenes were amplified in their respective color channel (e.g., 488nm for jGCaMP7b) and TH/5HT were labelled in the 647nm channel. After secondary antibody incubation, washing steps were repeated and the brain slices were then mounted onto microscope slides using mounting medium (Fluoromount; *Serva*, Germany) and coverslips (0.13-0.16mm thickness, 1871; *Carl Roth*, Germany).

#### Microscopy

Full overview images of immunostained brain slices were first acquired with an epifluorescence microscope (AxioObserver with Apotome 3; *Zeiss*, Germany) by stitching together individual tiles acquired with a 10× 0.45NA objective (Plan-Apochromat; *Zeiss*, Germany), controlled by the Zen software (v3.3; *Zeiss*, Germany). Detailed view of regions of interest within brain slices were then acquired with a confocal microscope (LSM 900 with Airyscan 2; *Zeiss*, Germany) using a 20× 0.8NA objective (Plan-Apochromat; *Zeiss*, Germany), controlled by the Zen software (v3.1; *Zeiss*, Germany). In both cases, presets corresponding to Alexa Fluor 488, 546 and 647nm were chosen. Intensity of excitation light, exposure time and gain of the photomultiplier tubes were manually adjusted to each sample in order to maximize signal quality while avoiding image saturation.

### Quantification and data analysis

#### Sample size

Sample size was determined according to past studies in the field and set to *n* = 6 animals for all experiments, with the exception of the joint LC-DR calcium recording experiment. In this case, with the goal of performing subgroup comparisons (by exchanging calcium indicators in LC-NA and DR-5HT), we increased the sample size to *n* = 4 animals × 2 subgroups = 8 animals. No differences were observed between pairs of indicators used, and data from all animals were thus pooled for following analyses.

#### Preprocessing of locomotion recordings

The position data was downsampled from 1800Hz to its nominal 20Hz acquisition rate. Speed was then computed as the first-order derivative on the cumulated position across treadmill laps. Locomotion bouts were defined as samples with instantaneous speed larger than 1cm·s^−1^, including samples 3s before and 8.5s after the motion event to account for the slow locomotion-induced pupil response. This conservative threshold allowed to remove locomotion bouts as well as small jolting movements which have been shown to also modulate brain activity^64^. To maximize the number of trials, the speed threshold for optogenetics experiments was set to 5cm·s^−1^.

#### Preprocessing of pupil recordings

Videos of the left eye were obtained at a 720 × 480-pixel resolution where 1mm corresponds to 120 pixels. Frames were manually cropped around the eye and downscaled to 256 pixels height. Eight markers equally spaced along the pupil edge (left, left dorsal, dorsal, dorsal right, right, right ventral, ventral left) were used to define the pupil. Pupil markers were localized on each individual eye frame using DeepLabCut^59^. In short, a pre-trained 50-layer convolutional neural network (ResNet-50) was further trained for 1.250.000 iterations on 1000 manually labeled eye frames from 20 videos using a dedicated GPU (Quadro P2000 5GB GDDR5; *Nvidia*, USA). An ellipse was fitted on coordinates of pupil markers with confidence ≥ 95% using a least squares method. Pupil size was estimated as the diameter of the fitted ellipse, which corresponds to the average of the ellipse’s semi-minor and -major diameters. When less than 5 pupil markers were available, the corresponding pupil size sample was marked as missing and was linearly interpolated from neighboring, available samples. A 5-sample wide median filter was then applied to the resulting pupil size trace in order to remove outlier samples. Finally, a 10Hz third-order Butterworth low-pass filter was applied to remove high-frequency noise.

#### Preprocessing of calcium recordings

First, a 5-sample wide median filter was applied to the calcium trace to remove outlier samples. Second, a 30Hz third-order Butterworth low-pass filter was applied to remove high-frequency noise. The decrease in baseline fluorescence caused by photobleaching was corrected by a 0.005Hz third-order Butterworth high-pass filter. The decrease in dynamic range resulting from photobleaching was furthermore corrected by linear scaling based on the 5% most extreme values estimated using a 60s-window running percentile. The baseline fluorescence (*F*_0_) was computed as the 8^th^ percentile of the photobleaching-corrected trace and used to estimate the relative change in fluorescence as: Δ*F*/*F*_0_ = (*F − F*_0_)/*F*_0_.

#### Additional common preprocessing

To ease downstream analyses, locomotion, pupil and brainstem activity were all resampled at 100Hz using linear interpolation. Furthermore, to allow for pooling of data across different sessions, all signals were z-scored separately for each session. Snapshots of calcium activity and pupil size were further low-pass filtered at 3Hz for illustration purposes.

#### Transient detection

To detect pupil events and calcium transients from spontaneous fluctuations (Fig. 1f, Fig. 3a), we computed the first-order derivative of the respective signal. In pupil recordings, we detected samples above/below the 90^th^/10^th^ percentile for pupil dilation/constriction events respectively, with a 400ms minimum duration between successive events. In calcium recordings, we detected samples above the 90^th^ percentile with a 500ms minimum duration between successive events.

#### Evoked responses

Pupil and calcium evoked responses were locked onto events of interest, epoched and baseline-corrected by regressing-out the effect of the baseline^128^ averaged in a time-window before event onset: from –5 to –3s for pupil/calcium transient analyses (Fig. 3a), and from –5 to 0s for responses to optogenetic activation (Fig. 5c, Fig. 6c, Fig. 8c).

#### Time-shifted analyses

For all analyses relying on time shifting such as cross-correlations, data samples at onset and offset of data bouts were discarded such that the resulting metric (e.g., correlation) was computed on the same data samples for all the different time lags considered. Brainstem-pupil lags varied from −10 to 10s (with 10ms increment).

#### Pupil phase analysis

Pupil phase-locked analyses (Fig. 2d and Fig. 3f) were performed by averaging brainstem activity into 129 phase bins obtained from the Hilbert transform of the 0.1-2Hz band-pass filtered signal of interest (pupil or LC-NA activity).

#### Identification of additive vs. multiplicative modulation

To assess whether LC-NA and DR-5HT pupil control were independent (Fig. 4c), we compared the explanatory power of two regression models:

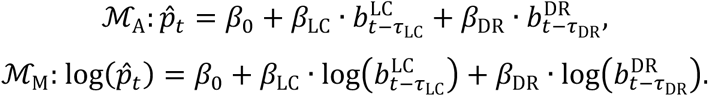

In the first model, LC-NA and DR-5HT activity levels (*b*) independently contribute to the expected fluctuations in pupil size (*p̂*). By contrast, in the second (log-linear) model, activity levels interact multiplicatively. Given that the two models have the same number of free parameters (3), the goodness-of-fit is simply quantified using the proportion of variance explained.

#### Stratified analysis

In this analysis, we assessed whether pupil size would still parametrically vary according to the activity of one brainstem system while the activity of the other brainstem activity would be at a particular given level (Fig. 4a-b). First, we aligned LC-NA and DR-5HT activities to pupil size based on the lag resulting in the largest correlation. The best lag was identified separately for each brainstem region and each mouse. Second, we arbitrarily defined 3 activity levels (–1.5, 1 and 3.5s.d.) and identified data samples for which the activity of the first brainstem region would fall into a small range around these values (±0.025s.d.). Third, and separately for each of these 3 activity levels, we grouped data samples based on the activity of the second brainstem region (13 bins from –4 to 6s.d. with a minimum of 10 data samples per bin). Thus, we could plot average pupil size conditionally on the activity of the first (3 levels) and second brainstem system (varying number of bins depending on the level considered). In another iteration of this analysis, we generalized the approach by computing the correlation between pupil size and the activity of the second brainstem system (without binning) while holding the activity of the first brainstem system along a finer grid (21 bins from –2 to 4s.d.).

#### Unique contribution to pupil size

To quantify the unique contribution of brainstem systems to fluctuations of pupil size (Fig. 4d-f), we adopted a double linear regression approach. First, we regressed-out the contribution of the first brainstem system (*b*_1_) to pupil size (*p*) to obtain pupil residuals (*p*′):

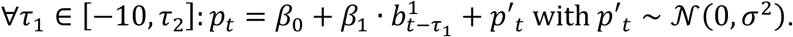

We then explained pupil residuals using the second brainstem system (*b*_2_):

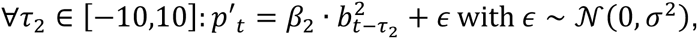

where *τ* denotes the brainstem-pupil lags and 𝒩 the normal distribution. The unique contribution is then defined as:

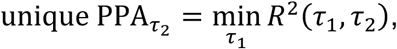

where *R*^2^ is the proportion of variance explained. The brainstem-pupil lag *τ*_2_ was varied between −10 and 10s with 100ms increment.

#### Pupil impulse response function

Based on past work^129–131^, we defined the pupil IRF (ℎ) as:

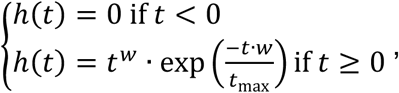

where *t* corresponds to time (in ms). The IRF has two free parameters: *w*, the width; and *t*_max_, the time-to-peak (in ms).

#### LC-based model of pupil response to DR activation

To obtain predictions of pupil size based on LC-NA activity in the joint DR-optogenetics/LC-recording experiment (Fig. 7), we convolved the continuous LC-NA activity trace by the pupil IRF defined above. Before fitting, pupil size and calcium activity were high-pass filtered at 0.01Hz to remove slow trends not captured by phasic changes in brainstem activity. Fitting of the two IRF parameters was performed on two held-out, opto-free sessions. The cost of cross-session generalization was assessed by fitting the IRF parameters on one opto-free session and assessing the proportion of variance explained in the other opto-free session. The fitted IRF was then used to convolve LC-NA activity recorded in sessions with optogenetic manipulations. Only offset and scaling parameters were adjusted separately in each optogenetic session. This LC-based model was finally compared to the observed pupil by locking its predictions onto the onset of DR-5HT optogenetic activation and by computing the proportion of explained variance over all sessions.

#### Model of pupil response to joint LC silencing and DR activation

To assess the independence between overlapping LC-NA- and DR-5HT-driven modulations of pupil size (Fig. 8d-f), we described the pupil response of joint manipulations (**p**_**LC**+**DR**_) as a linear weighted sum (with ***β*** > 0) of the pupil response to each optogenetic manipulation in isolation (**p**_**LC**_ and **p**_**DR**_) after alignment of light delivery onsets:

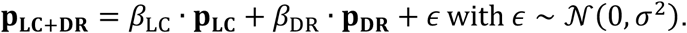

Only pupil response during optogenetic light (0 to 14s post orange light onset) was fitted.

#### Statistical testing

All figures show results averaged across mice. Unless stated otherwise, all statistical tests were two-tailed, with a type-1 error risk set at α = 0.05. For single- and two-group comparisons, non-parametric statistical testing was used: the Wilcoxon rank sum test for comparison of two independent samples, and the Wilcoxon signed rank test for comparison of paired conditions in single sample. For comparisons of more than two groups or conditions, ANOVAs were used.

#### Statistical testing for multi-dimensional data

To statistically compare multi-dimensional data (e.g., time-series, lag × frequency maps) while addressing the multiple comparison issue, we used non-parametric cluster-based statistical permutation testing^132^. We used the same type-1 error risk of α = 0.05 both at the sample and cluster levels and computed the *P*-value using all possible permutations of the experimental conditions for paired tests and 10,000 permutations for independent tests. For responses to optogenetic modulations, we ran the test only on samples post optogenetic stimulation onset.

#### Bootstrapping

Bootstrapping was used to estimate the null distribution of an evoked response. To do so, pseudo-onsets were randomly and uniformly sampled (with replacement) from all possible time labels in the recording. The same number of pseudo-onsets and true onsets were used. Epoching around pseudo-onsets and baseline correction was performed similarly as for the true onsets. This whole process was repeated 10,000 independent times per session and aggregated over sessions to yield the null distribution. We then used the 2.5 and 97.5% percentiles of this null distribution as thresholds.

## Supplementary figures

**Supplementary Fig. 1.**
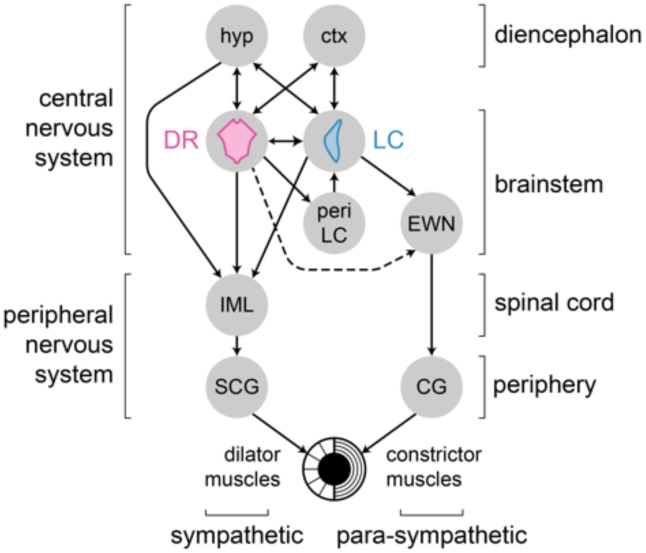
Summary of the main pupil control pathways. Pupil dilation and constriction are respectively caused by dilator and constrictor muscles of the iris, which are themselves respectively under the control of the sympathetic and para-sympathetic pathways^7,19,20,133^. The para-sympathetic pathway is composed of the ciliary ganglion (CG) which is under control of the Edinger-Westphal nucleus (EWN), itself receiving direct inputs from the LC. The LC also modulates the sympathetic pathway through direct input to the intermedio-lateral (IML) column of the spinal cord, itself connected to the superior-cervical ganglion (SCG). There also exists a hypothalamic relay mediating the modulation of IML by LC. Similarly, DR connects both directly and indirectly through the hypothalamus (hyp) to the IML column. In addition, DR connects to the peri-LC region^35^, where GABAergic interneurons are located and modulate LC^17^. Direct reciprocal connections between LC and DR have been identified^30–36^, with a stronger connection in the DR-to-LC direction. DR and LC also both receives inputs from the cortex (ctx), in particular from the frontal lobes^68,69^; an input which could synchronize their activities. Another candidate for a common input to LC and DR is the hypothalamus which entertains reciprocal connections with both^69^. The existence of a DR modulation of the para-sympathetic pathway via the Edinger-Westphal nucleus is debated^20^.

**Supplementary Fig. 2.**
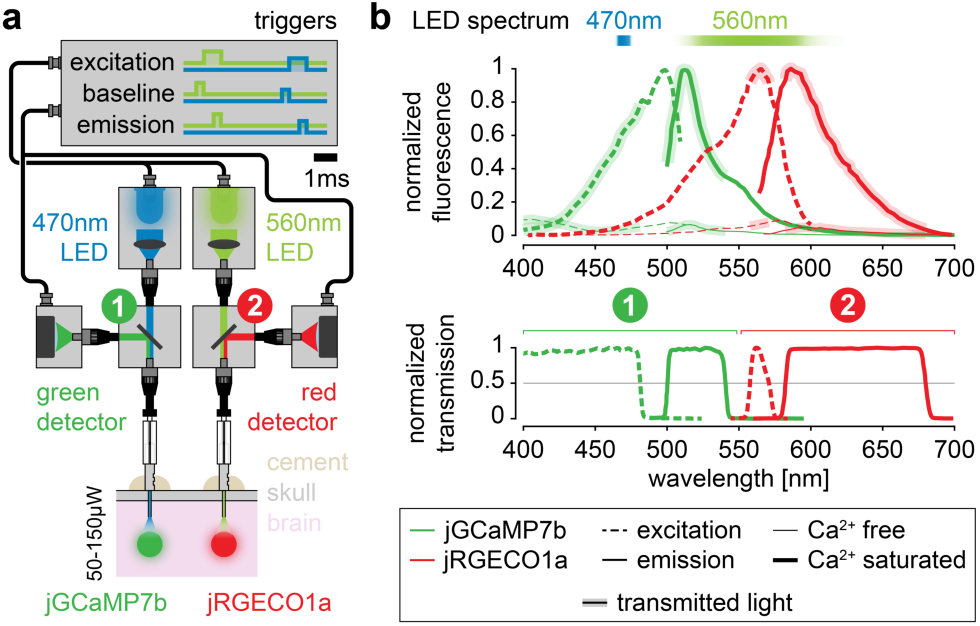
Cross-talk in population recordings is avoided by choice of filters and indicators. **a,** Apparatus for the dual-region, dual-color population recordings of calcium activity. The two excitation LEDs (470nm and 560nm) and baseline/emission measures are controlled by a micro-controller^127^ according to a time-division multiplexed acquisition scheme. Delivery of excitation light and measure of emitted fluorescence are performed through the same implanted fiber optic implant. Dichroic mirrors with incorporated band-pass filters collect emission light from the green (1: jGCaMP7b) and red (2: jRGECO1a) spectra using two distinct photodetectors. **b,** jGCaMP7b and jRGECO1a spectra are replotted from seminal publications^121,122^ (top plot) and transmitted light from the two dichroic mirrors are measured (bottom plot).

**Supplementary Fig. 3.**
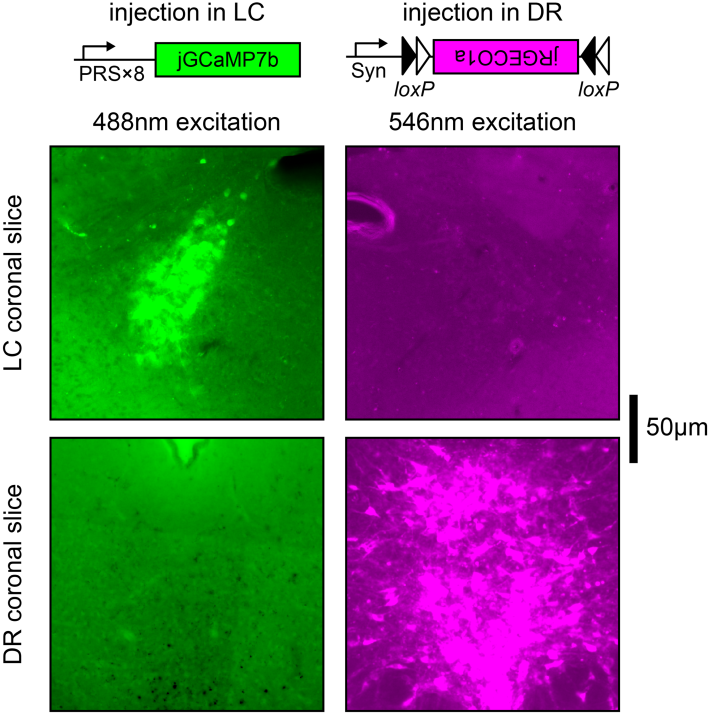
Absence of unspecific calcium indicator expression in LC and DR. Expression of genetically encoded calcium indicators (native fluorescence) in injected region (LC or DR) vs. the other region (DR or LC, respectively). Note that an identical scaling of the intensity distribution across pixels (which was saturated to reveal potentially weakly expressing cells in the non-injected region) was used for each injected vs. other region comparison (i.e., within in each column).

**Supplementary Fig. 4.**
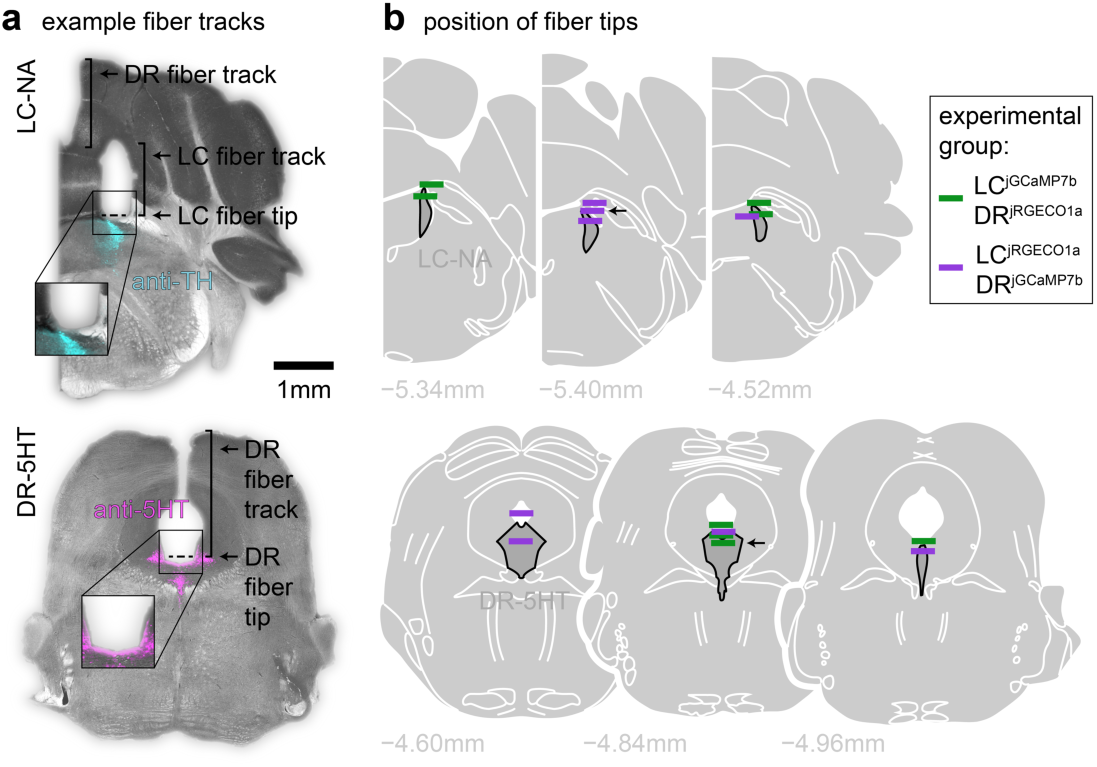
Position of implanted optic fibers near the targeted regions. **a,** Fiber tracks (400μm diameter optic fiber) and fiber tips highlighted on example coronal LC (top) and DR (bottom) slices. Targeted cells are colored and correspond to anti-TH (top) and anti-5HT (bottom) immunostaining. **b,** Fiber tips from all 8 mice mapped onto the Paxinos mouse atlas (after manual alignment with histology slices). Colors correspond to subgroups with different calcium indicators. Arrows correspond to the examples presented in **a**.

**Supplementary Fig. 5.**
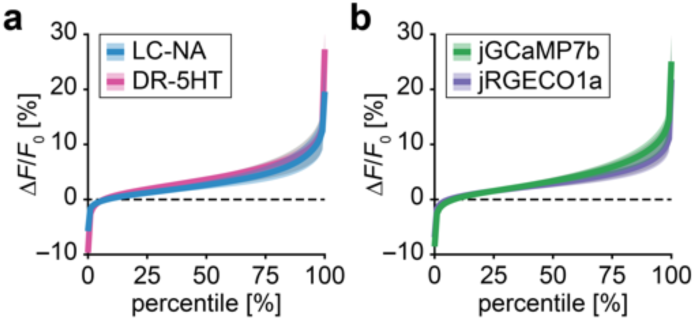
No effect of calcium indicator on signal quality. **a,** Mean ± s.e.m. distribution of signal strength between brainstem regions. Grey area: null distribution obtained after randomly shuffling mice across conditions (ɑ = 0.05; 10,000 permutations per session). **b,** Same as **a** but by splitting according to calcium indicator instead of brainstem region.

**Supplementary Fig. 6.**
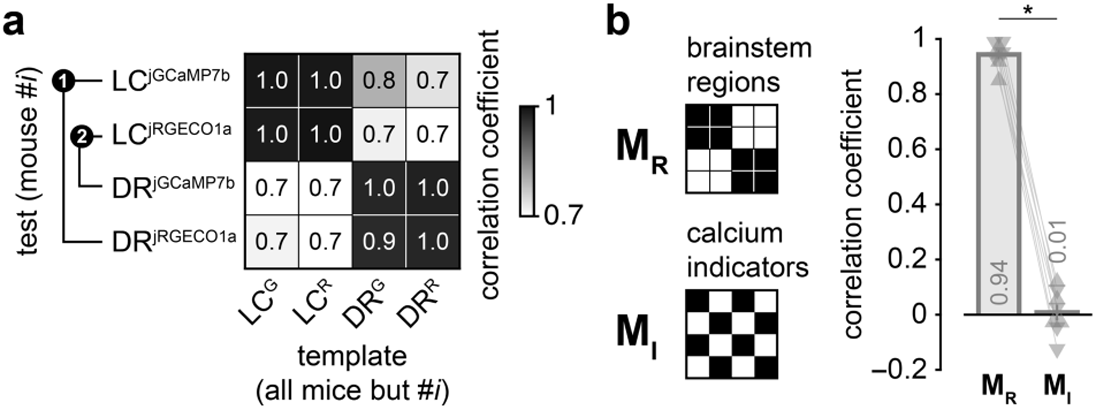
Similarity of calcium transients within brainstem region but not across calcium indicators. **a,** Mean correlation between activity transients across brainstem regions and calcium indicators. The two activity transients available from each mouse are compared against four template activity transients (pair #1 or #2) corresponding to all combinations of brainstem regions and calcium indicators averaged over all other mice. **b,** Mean ± s.e.m. correlation coefficients between observed (as in **a**) and theoretical correlation matrices describing the pure effect of brainstem systems vs. calcium indicators on activity transients. Upward-/downward pointing triangles: individual mice expressing LC^jGCaMP7b^-DR^jRGECO1a^/LC^jRGECO1a^-DR^jGCaMP7b^. Significant difference between correlation coefficients (Wilcoxon rank sum test, *P* < 0.05, *).

**Supplementary Fig. 7.**
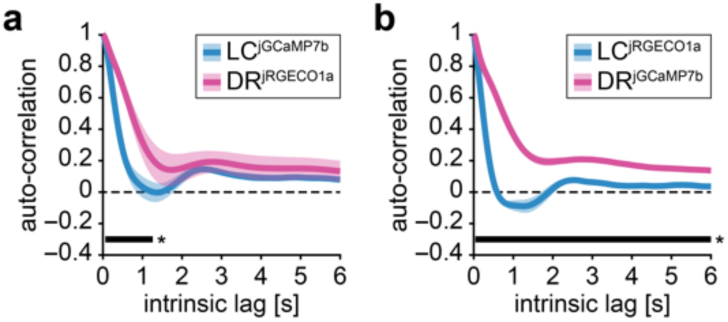
No effect of calcium indicator on population dynamics. **a,** Mean ± s.e.m. auto-correlograms separately for LC^jGCaMP7b^ vs. DR^jRGECO1a^. Horizontal black bars: significant difference between LC-NA and DR-5HT auto-correlograms (cluster-based permutation tests, *P* < 0.05, *). **b,** Same as **a** for LC^jRGECO1a^ vs. DR^jGCaMP7b^.

**Supplementary Fig. 8.**
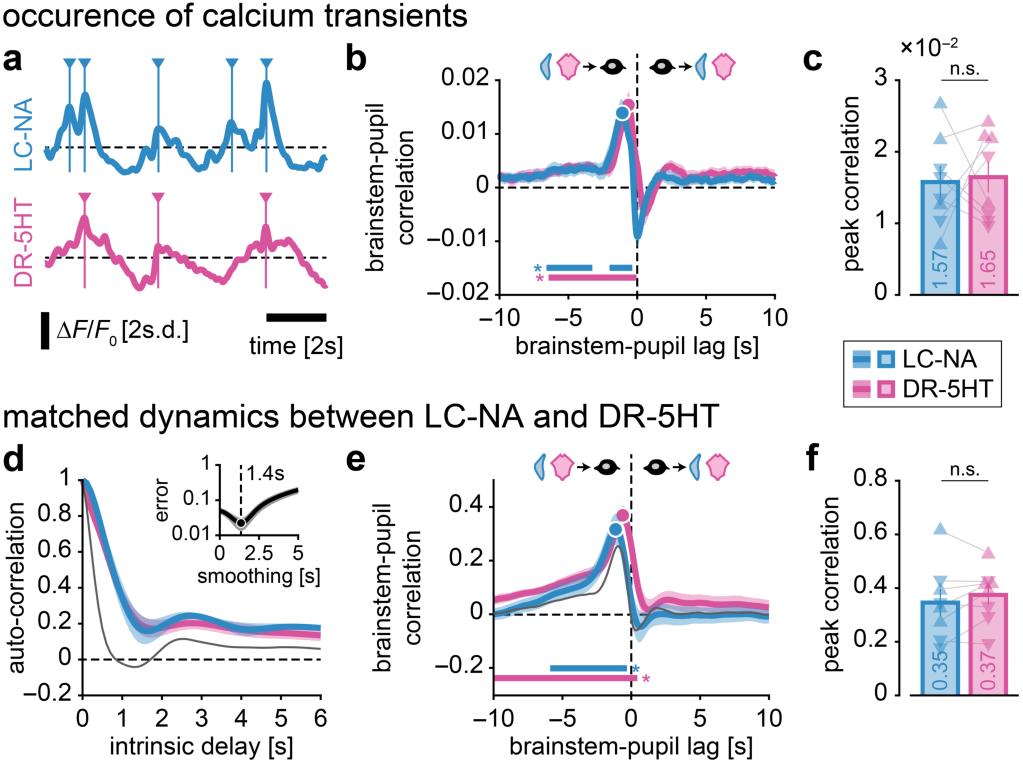
No difference in brainstem-pupil correlation after accounting for difference in population dynamics in LC-NA vs. DR-5HT. **a,** Activity peaks (local maxima) were identified from continuous fluctuations of LC-NA and DR-5HT activity. **b,** Mean ± s.e.m. brainstem-pupil cross-correlogram for LC-NA and DR-5HT after substituting continuous calcium fluctuations with calcium transient occurrence. Negative lags: brainstem leads pupil. Horizontal colored bars: significantly larger than zero correlation; no significant difference is found between brainstem regions (cluster-based permutation tests, *P* < 0.05, *). **c,** Mean ± s.e.m. peak cross-correlation (corresponding to dots in **b**). Upward-/downward-pointing triangles: individual mice expressing LC^jGCaMP7b^-DR^jRGECO1a^/LC^jRGECO1a^-DR^jGCaMP7b^. Non-significant (n.s.) difference in peak correlation (Wilcoxon rank sum test, *P* < 0.05). **d,** Mean ± s.e.m. auto-correlogram for DR-5HT activity (as in Fig. 1) and smoothed LC-NA activity. LC-NA activity was smoothed with a moving mean window until it matched the DR-5HT auto-correlation profile. Inset: mean absolute error between LC-NA and DR-5HT auto-correlograms across a range of smoothing window durations and the best smoothing window duration at the group level. Thin grey line: auto-correlogram for the raw LC-NA activity. **e,** Mean ± s.e.m. brainstem-pupil cross-correlogram for DR-5HT (as in Fig. 3) and LC-NA after auto-correlogram matching (same as **b**). **f,** Mean ± s.e.m. peak cross-correlation (corresponding to dots in **e**) like **c**.

**Supplementary Fig. 9.**
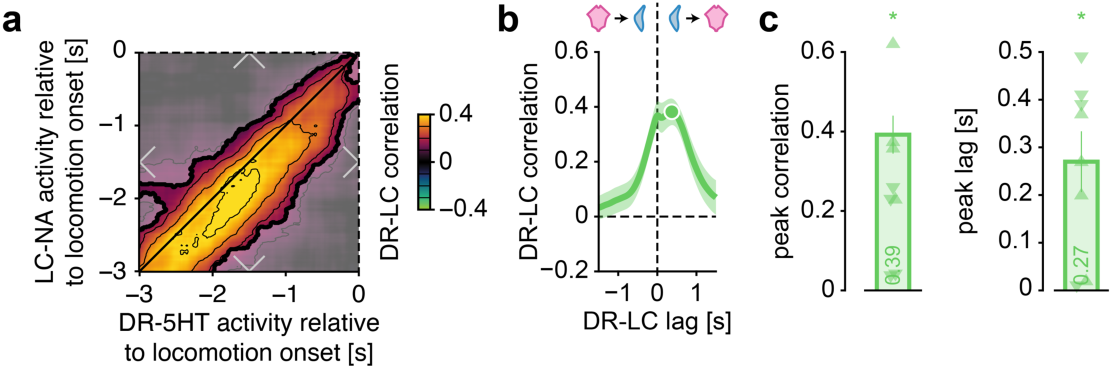
Brainstem-pupil coupling in intervals preceding locomotion. **a,** Mean correlation (across locomotion episodes) between LC-NA and DR-5HT relative to locomotion onset. Opaque regions: correlation significantly different from zero (cluster-based permutation tests, *P* < 0.05, *; n.s.: non-significant). **b,** Mean ± s.e.m. LC-DR cross-correlogram in periods preceding locomotion (corresponding to diagonal-centred square in **a**). **k**, Mean ± s.e.m. peak cross-correlation and corresponding LC-DR lag (dot in **b**).

**Supplementary Fig. 10.**
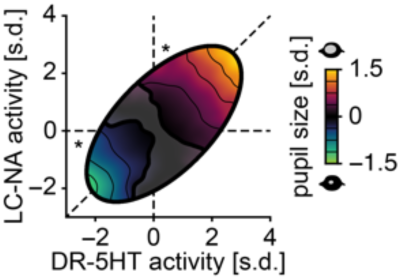
Cofluctuations between LC-NA, DR-5HT and pupil size. Mean pupil size as a function of a pair of activity levels of LC-NA and DR-5HT, while controlling for differences in timing and dynamics between the two brainstem systems. Opaque regions: brainstem activity significantly different from zero (cluster-based permutation tests, *P* < 0.05, *; n.s.: non-significant).

**Supplementary Fig. 11.**
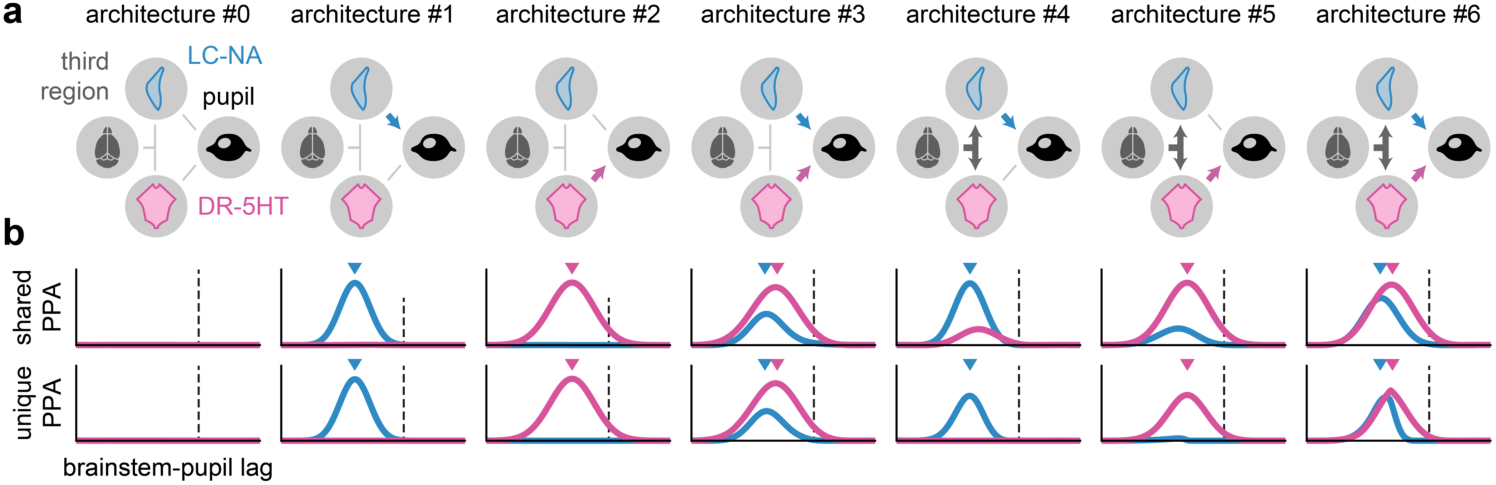
Estimation of unique pupil-predictive activity recovers simulated LC-DR-pupil architectures. **a,** Factorial exploration of LC-DR-pupil architectures. Pupil fluctuations were modeled as the linear, weighted sum of LC-NA and DR-5HT activities which could themselves be linearly modulated by a third region. Brain activity is modelled as auto-correlated random noise. Note that in this modelling framework, the cross-brainstem modulation is conceptually equivalent of the modulation of LC-NA and DR-5HT activity by the same third region (as in architectures #4-6), and is thus not shown here for the sake of simplicity. Architecture #0 is a control case where pupil is not modulated neither by LC-NA nor DR-5HT. **b,** Time course of shared (first row) and unique (second row) pupil-predictive activity (PPA) separately for LC-NA and DR-5HT. Downward-pointing triangles: the simulated peak of pupil-predictive activity. Simulations include slower dynamics for DR-5HT compared to LC-NA. Estimation of unique pupil-predictive activity accurately recovers the simulated architecture.

**Supplementary Fig. 12.**
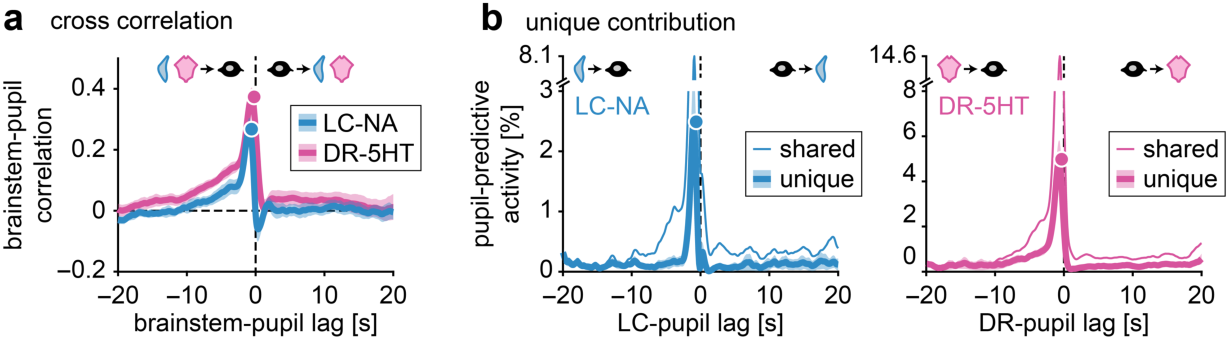
Control over entire DR time horizon in LC/DR independence analysis. **a**, Cross-correlation between brainstem activity and pupil size (as in Fig. 3c but for more protracted lags, here going from –20 to 20s). **b**, Unique and shared pupil-predictive activity from LC-NA and DR-5HT (as in Fig. 4f but after taking into account protracted lags into the analysis).

**Supplementary Fig. 13.**
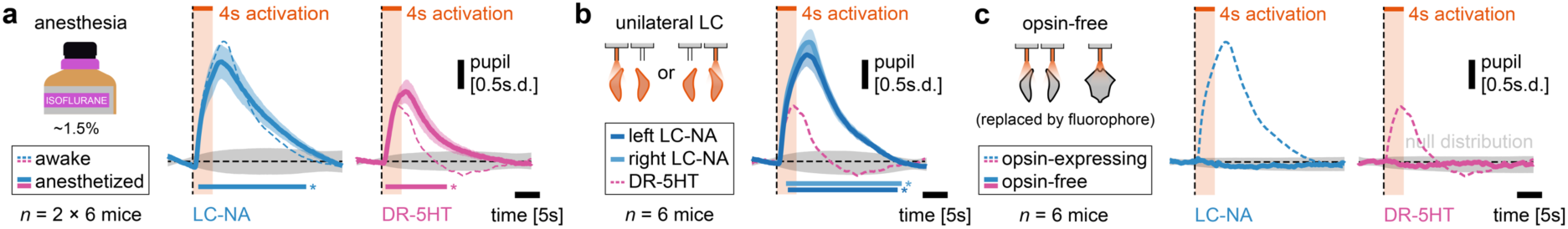
Control optogenetics experiments. **a,** Mean ± s.e.m. pupil response to optogenetic activation of LC-NA or DR-5HT in isoflurane-anesthetized (∼1.5%) animals. Horizontal bar: response significantly larger than zero (cluster-based permutation tests, *P* < 0.05, *). **b,** Mean ± s.e.m. pupil response to unilateral optogenetic activation of LC-NA. Horizontal bar: significant difference between unilateral LC-NA activation and DR-5HT activation (cluster-based permutation tests, *P* < 0.05, *). **c,** Mean ± s.e.m. pupil response to optogenetic activation of LC or DR in non-opsin expressing animals. Responses are not significantly different from zero (cluster-based permutation tests, *P* < 0.05).

**Supplementary Fig. 14.**
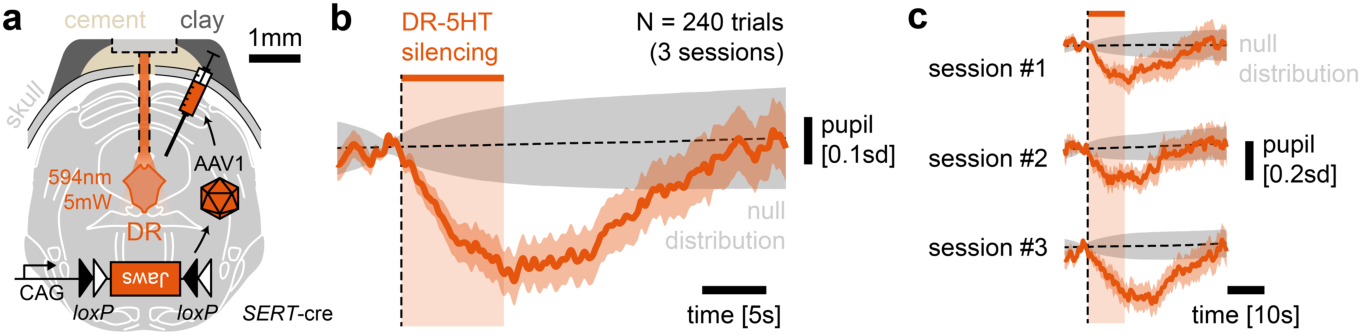
Pupil response to DR-5HT optogenetic silencing. **a**, The DR-5HT system of one mouse was optogenetically silenced using conditional expression of the silencing tools Jaws (similar conditions as in Fig. 8). **b**, Pupil response aligned to the onset of DR-5HT optogenetic silencing (8s of light delivery) reveals optogenetically-induced pupil constriction. **c**, Same as **b** but split by recording sessions on 3 consecutive days.

**Supplementary Fig. 15.**
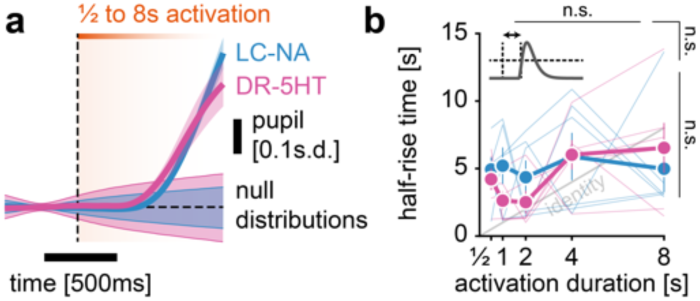
No difference in onset latency of pupil responses to optogenetic activation of LC-NA or DR-5HT. **a,** Mean ± s.e.m. pupil size in response to the optogenetic activation of LC-NA or DR-5H zoomed-in at light delivery onset (0 to 1s) demonstrates similar onset lag for LC-NA and DR-5HT. Shaded colored areas: the bootstrapped null distribution of evoked responses (ɑ = 0.05, 10,000 permutations per session). No significant difference between pupil-evoked responses is found (cluster-based permutation test, *P* < 0.05). **b,** Mean ± s.e.m. half-rise time of pupil responses to optogenetic stimulation, which corresponds to the time at which half the pupil peak is reached. Non-significant (n.s.) main effect of activation duration (top) brainstem region (right), and interaction between them (corner; ANOVA, *P* < 0.05).

**Supplementary Fig. 16.**
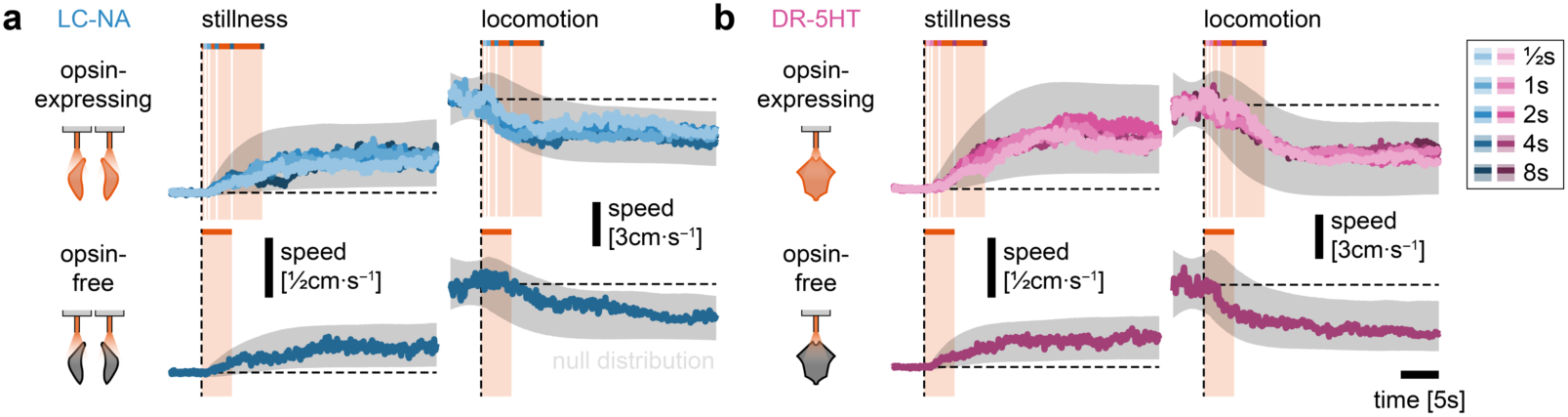
No initiation or interruption of locomotion upon optogenetic activation of LC-NA or DR-5HT. **a,** Mean ± s.e.m. speed in response to the optogenetic activation (½, 1, 2, 4, or 8s) of LC-NA in opsin-expressing (top row) and opsin-free (bottom row) animals, separately in trials in which the mouse was still (baseline speed < 5cm·s^−1^; left column) or moving (baseline speed > 5cm·s^−1^; right column). Shaded area: bootstrapped null distribution of evoked responses (ɑ = 0.05; 10,000 permutations per session). **b,** Same as **a** for DR-5HT optogenetic activation.

**Supplementary Fig. 17.**
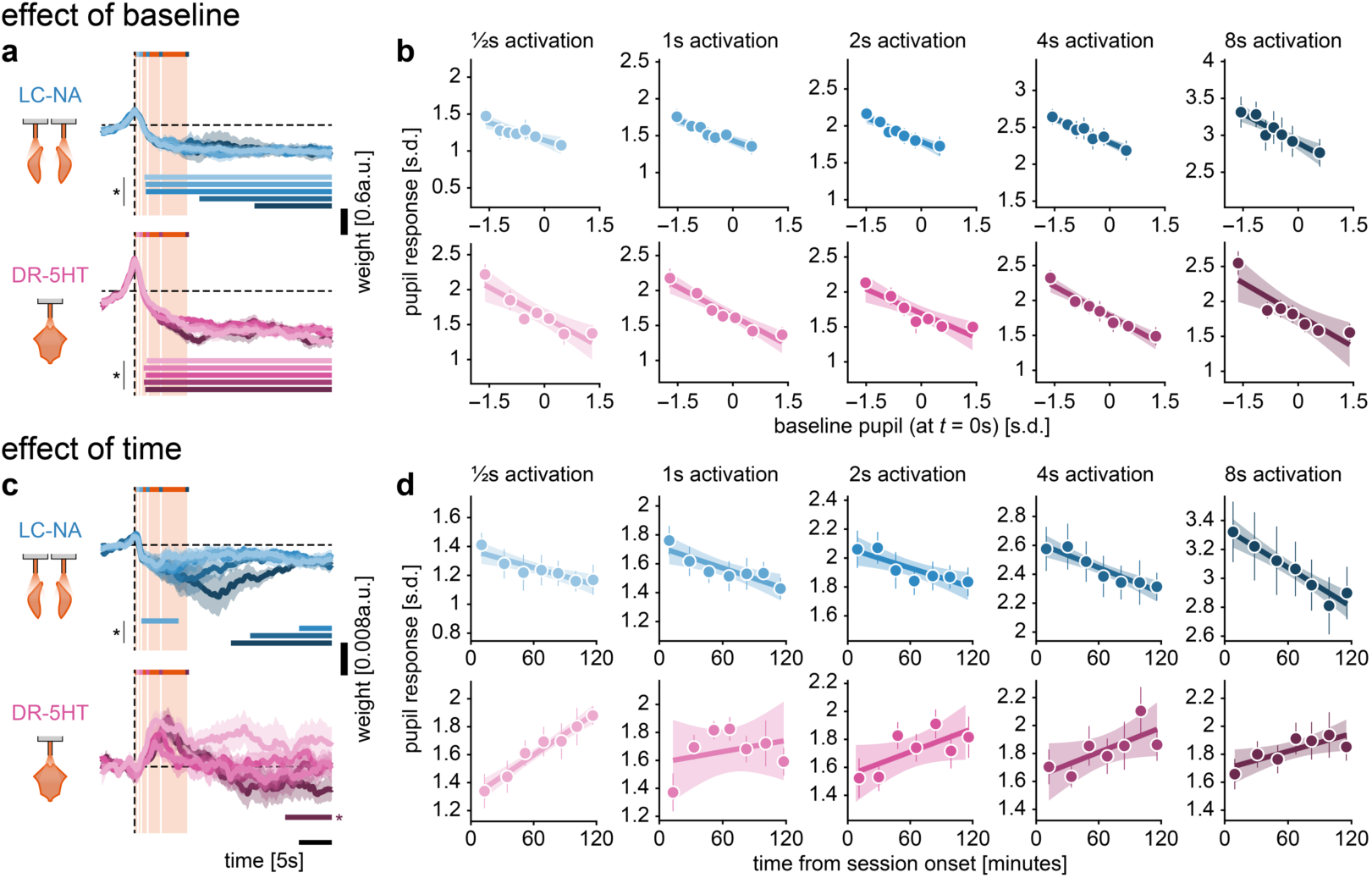
Modulation of pupil response to optogenetic activation by baseline pupil size and time. **a,** Mean ± s.e.m. time-course of the regression weight corresponding to the effect of baseline pupil on pupil size evoked by the optogenetic activation (½, 1, 2, 4, or 8s; color coded) of either LC-NA (first row) and DR-5HT (second row). Baseline pupil corresponds to pupil size at *t* = 0s with respect to optogenetic activation. In this analysis, the effect of time (see **c-d**) was regressed-out. Horizontal colored bars: weights significantly different from zero (cluster-based permutation tests, *P* < 0.05). **b,** Mean ± s.e.m. pupil response averaged in 7 bins of baseline pupil with linear fit ± 95% confidence intervals. The pupil response corresponds to the difference between the peak of pupil size post stimulation onset and the baseline pupil. **c,** Same as **a** but for the effect of time. Time corresponds to the onset of optogenetic activation in minutes, relative to session onset. In this analysis, the effect of baseline pupil (see **a-b**) was regressed-out. **d,** Same as **b** but for the effect of time.

**Supplementary Fig. 18.**
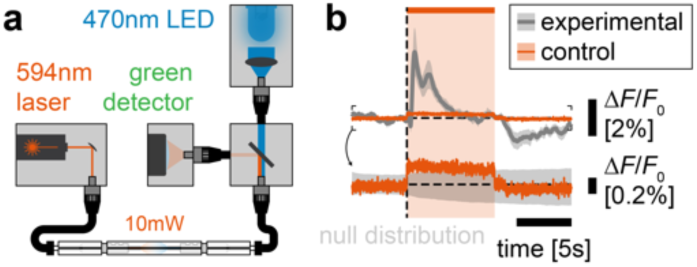
Negligible contamination of calcium recordings by optogenetic activation light. **a,** To quantify the amount of contamination from optogenetic stimulation light to calcium recordings, the stimulation and recording systems were connected without a mouse. Note that, contrary to real experiments, the optogenetic fiber optic implant was oriented directly towards the recording fiber optic implant. This approach thus gives a conservative upper bound on light contamination. Shaded area: bootstrapped null distribution of evoked responses (ɑ = 0.05; 10,000 permutations per session). **b,** Mean ± s.e.m. fluorescence in response to optogenetic light without a mouse (see **a**) and comparison with LC-NA activity. Optogenetic light was delivered for 8s, matching the longest stimulation trials used in real experiments. Optogenetic laser light was minimally picked up by the photodetector used for calcium recordings. Note that light contamination could occur through imperfect filtering of the dichroic mirror and/or a broad spectrum of the optogenetic activation light minimally extending to blue light.

**Supplementary Fig. 19.**
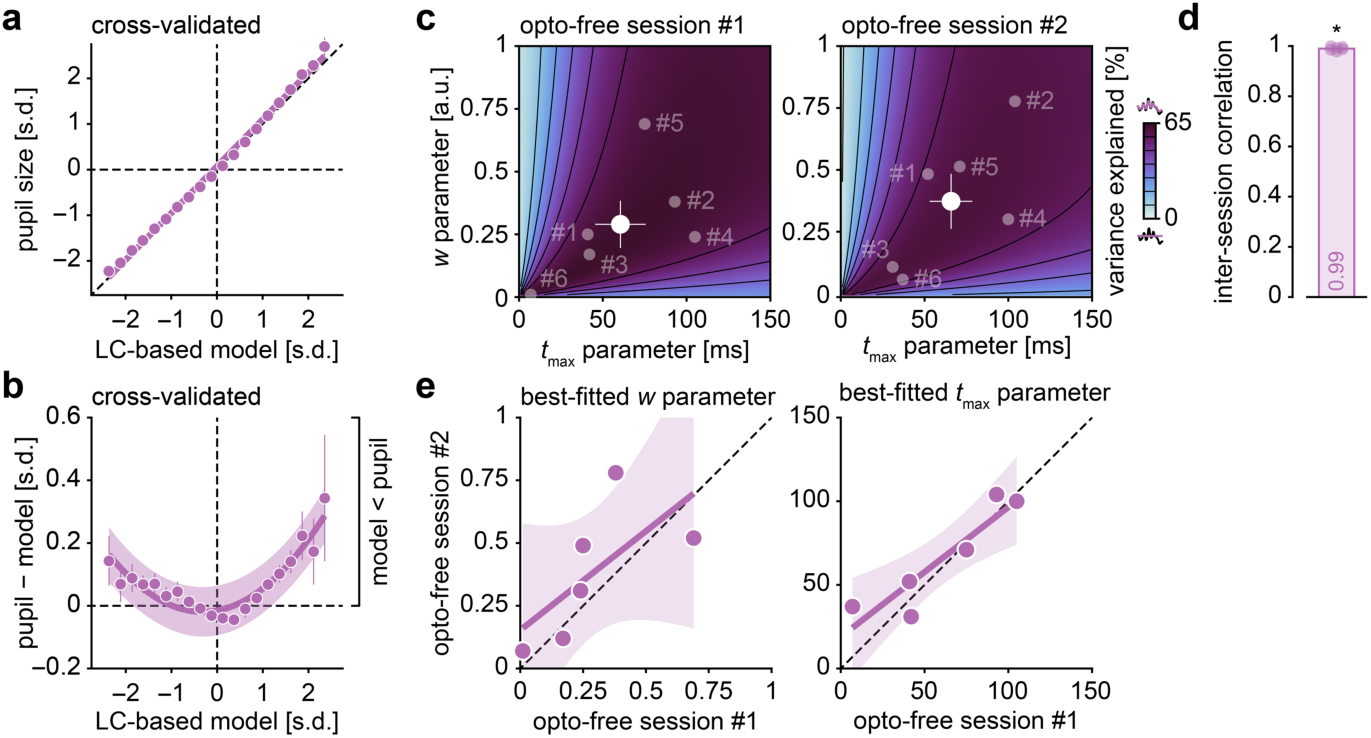
Agreement between IRF parameters fitted in separate opto-free sessions. **a,** Mean ± s.e.m. pupil size (in opto-session #*i*) as a function of the LC-based model (with IRF parameter fitted in opto-session #*j*) and linear fit ± 95% confidence interval demonstrate accurate model prediction throughout the whole pupil range. **b,** Mean ± s.e.m. difference between model predictions and pupil across a range of LC-based model predictions (same data as **a**) and quadratic fit ± 95% confidence interval demonstrate a slight underestimation of pupil size by the LC-based model for extreme pupil sizes. **c,** Mean proportion of variance explained and mean ± s.e.m. best-fitted IRF parameters (*w* and *t*_max_) in two separate opto-free sessions (left and right maps). Semi-transparent dots: individual mice. **d,** Mean ± s.e.m. correlation between proportion of variance explained across the parameter grid (in **c**). Semi-transparent dots: individual mice. Correlation significantly different from zero (Wilcoxon signed rank test, *P* < 0.05, *). **e,** Correlation between best-fitted parameters (left plot: *w*, right plot: *t*_max_) across mice in two opto-free sessions and linear fit ± 95% confidence interval.

**Supplementary Fig. 20.**
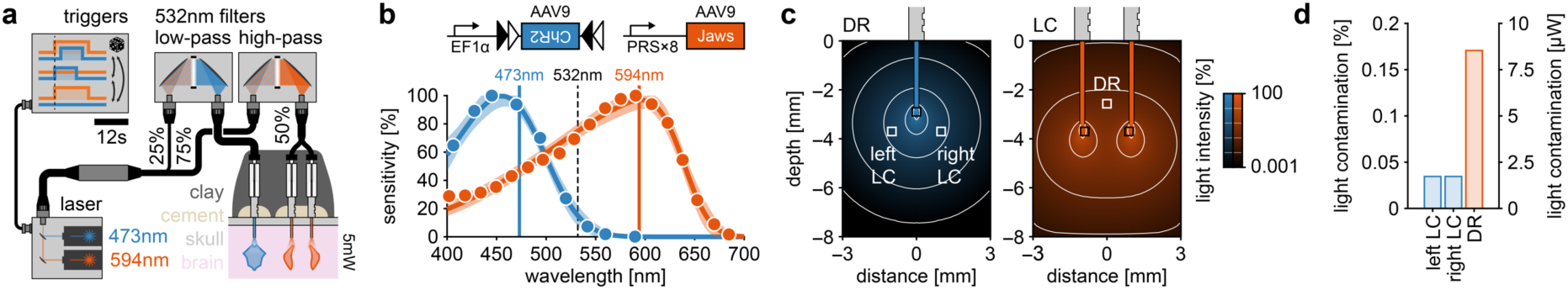
Optogenetic cross-activation is minimized by careful choice of filters, opsins and light intensity. **a,** Apparatus for the dual-region, dual-color optogenetic manipulation of LC-NA and DR-5HT. Cross-talk between LC and DR was minimized using a set of 532nm cutoff filters. Light delivery was calibrated to 5mW at the fiber tip. The three different trial (14s orange, 8s blue or 14s orange and 8s blue) types were pseudo-randomly interleaved. **b,** Spectral response profile of the optogenetic activation (ChR2) and silencing (Jaws) tools replotted from seminal publications^76,134^ and skewed gaussian fits ± 95% confidence intervals. **c,** Simulation of light spreading (5mW; using model from Johansson^135^ and code from Stujenske et al.^136^) from the tip of an fiber optic implant (200μm; 0.37NA) implanted either on top of DR or bilaterally on top of each LC (black squares). The relative position of the other brainstem region is projected in the 2D simulation (white squares). Note that fiber optic implants were implanted 100μm more lateral to the center of the LC to accommodate connection to two fiber-optic patch cords. **d,** Cross-region light contamination in estimated from simulations in **c** and according to calibrated light intensity at the fiber tip (5mW).

## Supplementary information

### Antibody list

Below is the immunostaining strategy for each brainstem region (LC and DR) in each experimental group.

| Group | LC transgene | DR transgene | Slice | Primary antibodies | Secondary antibodies |
| --- | --- | --- | --- | --- | --- |
| #1 | jGCaMP7b | jRGECO1a | LC | chicken anti-GFP<br>rabbit anti-TH | anti-chicken 488 (goat)<br>anti-rabbit 647 (goat) |
| #1 | jGCaMP7b | jRGECO1a | DR | rabbit anti-DsRed<br>goat anti-5HT | anti-rabbit 546 (donkey)<br>anti-goat 647 (donkey) |
| #2 | jRGECO1a | jGCaMP7b | LC | rabbit anti-DsRed<br>guinea pig anti-TH | anti-rabbit 546 (goat)<br>anti-guinea pig 647 (goat) |
| #2 | jRGECO1a | jGCaMP7b | DR | chicken anti-GFP<br>rabbit anti-5HT | anti-chicken 488 (goat)<br>anti-rabbit 647 (goat) |
| #3 | ChrimsonR-mRuby2 | — | LC | rabbit anti-TH | anti-rabbit 647 (goat) |
| #4 | jGCaMP7b | ChrimsonR-mRuby2 | LC | chicken anti-GFP<br>rabbit anti-TH | anti-chicken 488 (goat)<br>anti-rabbit 647 (goat) |
| #4 | jGCaMP7b | ChrimsonR-mRuby2 | DR | rabbit anti-5HT | anti-rabbit 647 (goat) |
| #5 | Jaws-tdTomato | ChR2-eYFP | LC | guinea pig anti-TH | anti-guinea pig 647 (goat) |
| #5 | Jaws-tdTomato | ChR2-eYFP | DR | chicken anti-GFP<br>rabbit anti-5HT | anti-chicken 488 (goat)<br>anti-rabbit 647 (goat) |

The technical specifications of each primary antibody are listed below:

| Species | Target | Company | Reference | Dilution |
| --- | --- | --- | --- | --- |
| chicken | GFP | Invitrogen | A10262 | 1:500 |
| rabbit | DsRed | TaKaRa | 632496 | 1:500 |
| rabbit | TH | Merck Millipore | AB152 | 1:500 |
| guinea pig | TH | Synaptic Systems | 213104 | 1:500 |
| rabbit | 5HT | Immunostar | 20080 | 1:500 |
| goat | 5HT | Immunostar | 20079 | 1:500 |

The technical specifications of each secondary antibody are listed below:

| Species | Target | Wavelength | Company | Reference | Dilution |
| --- | --- | --- | --- | --- | --- |
| goat | chicken | 488 | Invitrogen | A11039 | 1:1000 |
| goat | rabbit | 546 | Invitrogen | A11035 | 1:1000 |
| donkey | rabbit | 546 | Invitrogen | A10040 | 1:1000 |
| goat | rabbit | 647 | Invitrogen | A27040 | 1:1000 |
| donkey | goat | 647 | Invitrogen | A21447 | 1:1000 |
| goat | guinea pig | 647 | Invitrogen | A21450 | 1:1000 |

### Comparison of calcium transients across indicators and regions

Two calcium transients (*T*) are obtained per mouse: one for each brainstem region (*R*), recorded using a particular calcium indicator (*I*). For each mouse, the time-course of the two calcium transients were correlated against the two corresponding calcium transients averaged over all other mice (*T̄*). A 2 × 4 matrix (**M**) was then created by aggregating correlation coefficients together (*r*):

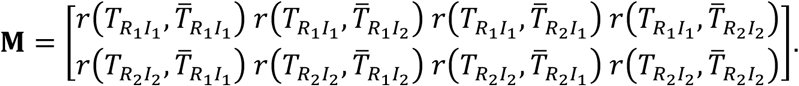

Finally, this empirical matrix (Supplementary Fig. 6a) was correlated separately against two theoretical matrices describing the pure effect of brainstem regions (**M_R_**) or calcium indicators (**M_I_**):

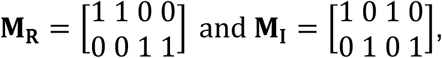

to quantify the similarity between calcium transients obtained from different brainstem regions (LC-NA vs. DR-5HT) vs. from different calcium indicators (jGCaMP7b vs. jRGECO1a; Supplementary Fig. 6b).

### Simulation of optogenetic light propagation

To simulate the amount of light propagating from one brainstem region to the other in the LC-silencing/DR-activation experiment (Supplementary Fig. 20c-d), we used Monte Carlo simulation of a random walk of photon packets (100 packets of 100,000 photons) through space (10μm precision) using a model^135^ and code implementation^136^ previously proposed. Simulated fiber optic implant had the same optical properties (200μm diameter, 0.22NA) as those employed in main experiments and simulated lights of the same wavelengths were used (473nm and 594nm). In LC simulations, we used two fiber optic implants. Position of the second brainstem region from which to measure light contamination was projected onto the 2D simulations according to LC-DR distance.

